# Sex-based difference in immune responses and efficacy of the pneumococcal conjugate vaccine

**DOI:** 10.1101/2024.04.15.589579

**Authors:** Essi Y. I. Tchalla, Anagha Betadpur, Andrew Y. Khalil, Manmeet Bhalla, Elsa N. Bou Ghanem

**Author notes:** Address correspondence to: Elsa N. Bou Ghanem, 955 Main Street, Buffalo, NY, 14203. Co-first authors (Essi Y. I. Tchalla, Anagha Betadpur).

## Abstract

Vaccine-mediated protection and susceptibility to *Streptococcus pneumoniae* (pneumococcus) infections are influenced by biological sex. The incidence of invasive pneumococcal disease remains higher in males compared to females even after the introduction of the pneumococcal conjugate vaccine (PCV). However, sex-based differences in the immune response to this conjugate vaccine remain unexplored. To investigate those differences, we vaccinated adult male and female mice with PCV and assessed cellular and humoral immune responses. Compared to females, male mice displayed lower levels of T follicular helper cells, germinal center B cells and plasmablasts, which are all required for antibody production following vaccination. This was linked to lower IgG and IgM levels against pneumococci and lower isotype switching to IgG3 in vaccinated males. Additionally, sera of vaccinated male mice had lower efficacy in several anti-pneumococcal functions including neutralization of bacterial binding to pulmonary epithelial cells as well as direct cytotoxicity against *S. pneumoniae*. Importantly, while the vaccine was highly protective in females, vaccinated males succumbed to infection more readily and were more susceptible to both lung-localized infection and systemic spread following *S. pneumoniae* challenge. These findings identify sex-based differences in immune responses to PCV that can inform future vaccine strategies.

## Introduction

Susceptibility to infections and the degree of protectivity by vaccines differ among people and populations and one determinant that can influence these differences is biological sex (1–3). Studies have shown that males are generally more susceptible to infectious diseases compared to females (1–4). Additionally, females tend to have higher immune responses to vaccines, as has been shown in the case of several vaccines including influenza, yellow fever, hepatitis B, smallpox, and diphtheria (1, 5–7). Although various studies have shown the differences in antibody responses between males and females, the mechanisms behind these sex-based differences are still understudied.

In response to vaccination, antibodies can be produced in a T cell dependent or T cell independent manner depending on the type of vaccine (8, 9). In the case of T-independent vaccines (such as purified polysaccharide vaccines), antibodies are typically produced by marginal zone B cells in secondary lymphoid organs while antibodies to T-dependent vaccines (such as polysaccharide-protein conjugate vaccines) are typically produced by germinal center (GC) B cells with the help of T follicular helper cells (Tfhs) (8–10). For T cell-dependent vaccines, Tfhs induce GC B cells to undergo differentiation into antibody secreting cells as well as class switch recombination and somatic hypermutation to produce antibodies specific to the antigen (11–13). As a result, different isotypes of antibodies are produced with high affinity to the antigen which is important for their functionality (14). Antibodies can confer protection to the host through various functions including neutralization, complement-mediated lysis, antibody-dependent cellular cytotoxicity, and/or phagocytosis of the pathogen (14). How sex-based differences affect antibody function against infectious diseases is not well understood.

*Streptococcus pneumoniae* (pneumococcus) are a group of encapsulated Gram-positive bacteria categorized by serotypes, based on composition of the polysaccharides in the capsule, more than 100 of which have been identified so far (15). These bacteria can cause pneumonia as well as invasive pneumococcal disease (IPD) including otitis media, meningitis and bacteremia leading to infection of distal organs such as the heart (16, 17). Pneumococcal infections, especially community-acquired pneumonia remain a major cause of mortality and morbidity despite the availability of 2 vaccine formulations against the pneumococcus (10, 18). The pneumococcal polysaccharide vaccine (PPSV) is composed of purified pneumococcal polysaccharides that can directly activate antibody production by B cells and is currently recommended for use in people aged ≥65 years old (9, 19). The pneumococcal conjugate vaccine (PCV) is composed of purified polysaccharides that are linked to a carrier protein (CRM197) and requires T cell help for antibody production by B cells (10). PCV is currently recommended for routine use in children aged ≤2years, individuals at risk, and people ≥65 years old (19).

Sex-based differences in the incidence of pneumococcal diseases have been reported, with males across different age groups being more susceptible than their aged-matched female counterparts (20–25). Additionally, sex-based differences in immune response to PPSV have also been reported in adults, with age influencing those differences (26–29). Studies have reported conflicting results with some showing more IgG in males versus others finding more in females (28, 30). However, studies have consistently found higher incidence of community-acquired pneumonia in PPSV-vaccinated aged males, suggesting a decline in antibody function (26, 27, 29). While studies investigating sex-based differences in immune response to PCV are fewer, population surveillance has shown that after the introduction of PCV to the population, the rates of invasive pneumococcal disease were still higher in males compared to females across age groups (20, 24). This suggests that females are better protected by PCV compared to males, however, studies looking at both cellular and humoral responses to PCV to understand the mechanisms behind the sex-based differences in vaccine protection are needed.

In this study, using a preclinical murine model, we evaluated sex-based differences in immune responses to PCV and found that both humoral and cellular responses following vaccination were lower in males compared to females. The lower antibody production in males as compared to females correlated with lower protection against subsequent pneumococcal infection. This study highlights sex-based differences in immune responses to PCV and broadens our understanding of the sex-based differences in vaccine efficacy which could inform future decisions on vaccine administration and design.

## Methods

### Mice

For all experiments, adult (12 weeks old) male and female C57BL/6 mice were purchased from Jackson Laboratories (Bar Harbor, Maine). Mice were housed in a specific-pathogen-free environment at the University at Buffalo, rested for two weeks and then used in experiments in accordance with Institutional Animal Care and Use Committee guidelines.

### Immunization and sera collection

Mice were injected intramuscularly with 50ul of the Polysaccharide Conjugate Vaccine (PCV) Prevnar 13 (Wyeth Pharmaceuticals) or Phosphate Buffered Saline (PBS) into the caudal thigh muscle. This single dose intramuscular injection models what is recommended for adults receiving PCV (31). 10ul of blood from each mouse was collected from the tail vein at weeks 0, 2 and 4 following vaccination in 90 μl of anticoagulant (75 μl of 1 M sodium citrate, 50 μl of 1 M citric acid, and 12.375 ml of PBS pH) and saved at −80°C for quantification of antibody titers. Sera from each mouse across three separate experiments were collected via cardiac puncture 4 weeks following vaccination, pooled per group, aliquoted and saved at −80°C for quantification of antibody functions in subsequent assays.

### Bacteria

*Streptococcus pneumoniae* TIGR4 strain (serotype 4) and EF3030 strain (serotype 19F) were a kind gift from Andrew Camilli and Anthony Campagnari respectively. Bacteria were grown to mid-exponential phase in Todd–Hewitt broth (BD Biosciences) supplemented with Oxyrase at 37°C/5% carbon dioxide. Aliquots were frozen at −80°C in growth media supplemented with 20% glycerol. Prior to use, bacteria were thawed on ice, washed, and diluted in PBS. Bacterial enumerations were confirmed by serial dilution and dribbled plating on Tryptic Soy Agar (TSA) plates supplemented with 5% sheep blood (Northeast Laboratory Services).

### Flow cytometry for T and B cell assessment

On days 7, 14 and 21 following immunization, mice were euthanized and spleens and vaccine draining lymph nodes (vLNs-inguinal and popliteal) were harvested. Single cell suspension of splenocytes and vLNs cells were prepared in RPMI 1640 media supplemented with 10% Fetal Bovine Serum by mashing the organs. Following red blood cells lysis with an in-house hypotonic solution (8.29g ammonium chloride, 1g sodium bicarbonate, 0.038g trypsin/ethylenediaminetetraacetic acid (EDTA) and 1L of water), the remaining cells were enumerated using trypan blue staining and a hematocytometer. In separate staining panels, 2×10^6^ cells per well were incubated with Fc block (2.4G2, BD) and Live/Dead dye (Invitrogen) and stained with CD4 (RM4-5, BD), TCRβ (H57-597, BD), CD11a (M17/4, BD), CD49d (R1-2, BD), PD1 (J43, Invitrogen), CXCR5 (SPRCL5, Invitrogen), ICOS (76.17G9, Invitrogen), CD40L (MR1, BD), CD45R/B220 (RA3-6B2, BD), CD138 (281-2, BD), CD95 (SA367H8, Biolegend), and/or CD38 (90, Biolegend) antibodies. Following a 30-minute incubation on ice in the dark, cells were washed with FACS Buffer and incubated in Fix/Perm buffer from the FOXP3/Transcription factor staining buffer set (eBioscience) overnight. The next day, cells were washed in Perm buffer and stained in separate panels with FOXP3 (FJK-16s, Invitrogen), RORγT (931-378, BD), Tbet (eBio4B10, eBioscience) and GATA3 (L580-823, BD). Following a 45 mins incubation on ice in the dark, cells were washed in Perm buffer, resuspended in FACS buffer (HBSS, 1% FBS and 0.1% sodium azide) and fluorescence was assessed using a BD Fortessa. At least 20,000 cells in the CD4^+^TCRβ^+^ or B220^+^ respectively were collected and analyzed using the FlowJo software.

### Antibody Enzyme-Linked Immunosorbent Assay

Antibodies in sera were assessed using an ELISA assay as previously described (12, 32, 33). Briefly, Nunc-maxisorp plates (Thermo Scientific) were coated at 4°C overnight with 2×10^5^ colony forming unit (CFU) per well of heat killed *Streptococcus pneumoniae* serotype 4. The next day, plates were washed in PBS supplemented with 25% Tween (PBS/T) and sera were added to the plates. After a 2-hour incubation, plates were washed and incubated with diluted horseradish peroxidase (HRP) conjugated anti-mouse IgM (Invitrogen), IgG (Millipore Sigma) or IgG1, IgG2b, IgG2c or IgG3 (Southern Biotech). One hour later, plates were washed, TMB substrate (Thermo Scientific) was added, and absorbance was read at 650nm using a Biotek microplate reader for 10 minutes at 1-minute intervals. Antibody units were calculated as a percentage of standard sera included in each assay. Standard sera were obtained from mice that were intranasally inoculated with *S. pneumoniae* serotype 4 weekly for 3 weeks and immunized with PCV at week 4. Four weeks following vaccination, sera were collected from the mice, aliquoted and stored at −80°C. Antibody units were then calculated in μg/ml from the standard sera which was quantified using antibody quantification kits from Invitrogen for IgG (88–50400) and IgM (88–50470).

### Cytokine/Chemokine Array

On days 7 and 14 following immunization, mice were euthanized, and spleens were harvested. Single cell suspension of splenocytes were prepared in 2ml of RPMI 1640 media supplemented with 10% Fetal Bovine Serum by mashing the organs and saved at −80°C. Samples were thawed on ice prior to use, lysed with Triton-X (VWR, Cat# 0694) and quantified for protein concentrations using the Pierce BCA protein assay kit (Thermofisher, Cat# A55864) per the manufacturer’s instructions. All samples were brought to the same protein concentrations and used for pan-cytokine assessment using the Proteome Profiler Mouse XL Cytokine Array kit (R&D, Cat# ARY028) following the manufacturer’s instructions. Pan-cytokine protein membranes were read using the BioRad ChemiDoc MP Imaging System. The intensity for each dot on the blots was quantified using the Quick Spots Image Analysis Software and data exported to Excel. The intensity values of each blot were blanked using their respective negative controls (buffer only). Intensity values across multiple blots were normalized using the blot with the highest reference points (Standard). To present the data, fold change of the values from vaccinated PMN-depleted mice over those of vaccinated PMN-sufficient mice were calculated and log transformed.

### Antibody binding to the surface of *S. pneumoniae*

To measure the binding of antibodies to the surface of *S. pneumoniae*, sera collected 4-weeks post vaccination were incubated with GFP-expressing pneumococcus (TIGR4) at 1-, 3- and 6% of total assay volume in HBSS (with Ca^2+^ and Mg^2+^). Following a 30-minute incubation at 37°C, cells were washed with FACS buffer and labeled with an APC-tagged anti-mouse IgG (polyclonal, APC) antibody for 30 minutes on ice in the dark. Cells were then washed, resuspended in FACS buffer and fluorescence was assessed using a BD Fortessa. At least 10,000 cells were analyzed using the FlowJo software.

### Antibody neutralization of *S. pneumoniae* from binding to epithelial cells

The ability of antibodies to neutralize binding of the pneumococcus to epithelial cells was measured using H292 human pulmonary mucoepidermoid carcinoma derived cells (ATCC) as previously described (32, 34). Briefly, 2.5×10^5^ epithelial cells were seeded in tissue culture-treated flat bottom 96-well plates (Corning) and adhered overnight. The next day, the cells were washed three times with PBS and infected with the *S. pneumoniae* TIGR4 strain at a multiplicity of infection (MOI) of 10 in antibiotic-free media pre-opsonized with different sera. Sera collected 4-weeks post vaccination were incubated with TIGR4 at 10- and 20% of total assay volume in HBSS (with Ca^2+^ and Mg^2+^). In prior work we found that at least 10% sera were required to neutralize bacterial binding to the pulmonary epithelium and no neutralization is observed with lower concentrations even with standard sera from hyperimmune mice (32). After a 30-minute incubation at 37°C, the sera/bacteria mix was added to H292 cells, the plates spun down to facilitate host-bacterial interaction and incubated for 1 hr at 37°C/CO_2_. The cells were then gently washed 5 times with PBS to remove unbound bacteria, lifted with 0.05% EDTA (Invitrogen), and mixed well to produce a homogeneous solution. Dilutions were then plated on blood agar plates to determine CFU. The percentage of bound bacteria was determined with respect to a no cell control where bacteria were added to the wells and incubated for an hour with the same sera conditions. The number of bacteria bound to H292 cells in the naïve sera condition for each sex was set as 100% and relative changes in bacterial binding were then calculated for the immune sera conditions.

### Opsonophagocytic Killing Assay (OPH)

The ability of sera to promote opsonophagocytic killing of *S. pneumoniae* by neutrophils was assessed using a Opsonophagocytic Killing Assay (OPH) assay as previously described (12, 32, 35). Briefly, neutrophils were isolated from the bone marrow of mice using density centrifugation with Histopaque 1119 and Histopaque 1077 (Sigma). Neutrophils were resuspended in HBSS (without Ca^2+^ or Mg^2+^) media supplemented with 0.1% gelatin and used in the assay. Sera collected 4-weeks post vaccination were incubated with TIGR4 at 1-, 3- and 6% of total assay volume for 5 minutes and then added to neutrophils in HBSS (with Ca^2+^ and Mg^2+^). Following a 40-minute incubation at 37°C, reactions were dribbled plated on TSA plates with 5% sheep blood. Percent killing was calculated relative to a no neutrophils control for each sera condition.

### Opsonophagocytic uptake flow cytometry assay

The ability of sera to promote opsonophagocytic uptake of *S. pneumoniae* by neutrophils was assessed via flow cytometry as previously described (36). Briefly, neutrophils were isolated from the bone marrow of mice using density centrifugation as described above. Sera collected 4-weeks post vaccination were incubated with GFP-tagged TIGR4 at 1-, 3- and 6% of total assay volume for 30 minutes and then added to neutrophils in HBSS (with Ca^2+^ and Mg^2+^). Following a 5-minute incubation at 37°C, cells were stained with Fc block (2.4G2, BD) and anti-pneumococcal serotype 4 rabbit serum (Statens Serum institutes) for 30 minutes at room temperature. Afterwards, cells were stained with a secondary PE-tagged anti-rabbit IgG antibody for 30 minutes on ice in the dark. Cells were then washed, resuspended in FACS buffer and fluorescence was assessed using a BD Fortessa. At least 10,000 cells were analyzed using the FlowJo software.

### Complement binding the surface of *S. pneumoniae*

To measure the binding of complement to the surface of *S. pneumoniae*, sera collected 4-weeks post vaccination were incubated with the pneumococcus (TIGR4) at 1-, 3- and 6% of total assay volume in HBSS (with Ca^2+^ and Mg^2+^). Following a 30-minute incubation at 37°C, cells were washed with FACS buffer and labeled with a FITC-tagged anti-mouse C3 antibody (Polyclonal, ICL) for 15 minutes on ice in the dark. Cells were then washed, resuspended in FACS buffer and fluorescence was assessed using a BD Fortessa. At least 10,000 cells were analyzed using the FlowJo software. Heat-inactivated sera were used as negative controls and showed no C3 binding.

### Animal Infections

Mice were infected intratracheally with 10^7^ CFU of *S. pneumoniae* serotype 4 (TIGR4) or serotype 19F (EF3030) using the tongue pull method as previously described (12, 37). Twenty-four hours following infection, mice were euthanized for enumeration of bacterial burden. Briefly, lung and blood were collected, homogenized, serially diluted and dribbled plated on TSA plates with 5% sheep blood. A separate set of mice (for TIGR4 only) was monitored for 7 days post infection. Mice were blindly assessed for clinical signs of disease progression including breathing, posture, weight, activity, grooming, and respiration ranging from 0 (healthy) to 21 (severely sick) as previously described (12, 32, 35, 37).

### Statistical Analysis

All statistical analysis was done using GraphPad Prism version 9 software. Data were checked for normality distribution using the Shapiro-Wilk test prior to statistical analysis. Significant differences were determined by Fisher exact test, 1-way analysis of variance followed Šídák’s or Dunnett’s multiple comparisons test, Kruskal-Wallis 1-way analysis of variance followed Dunn’s multiple comparisons, or 1-sample t test. Survival was analyzed using the log-rank (Mantel-Cox) test. All *p* values <0.05 were considered significant.

## Results

### Sex-based differences in cellular immune responses in the vaccine-draining lymph nodes following PCV vaccination

Immune response to conjugate vaccines involves both B and T cells (38). Germinal center (GC) B cells and T follicular helper cells (Tfh) interact in a contact dependent manner to induce B cell differentiation and antibody production (38–40). Therefore, on days 7, 14 and 21 post PCV vaccination, we assessed T and B cell responses in the spleen and vaccine draining lymph nodes (vLNs) of 12 weeks old adult male and female mice. We first looked at Tfhs (CXCR5^+^PD1^+^CD4^+^TCRβ^+^ T cells) (gating strategy in Fig. S1). Upon vaccination, there was a significant increase in the number of Tfhs in vLNs of female but not male mice, with a peak at 14 days post vaccination (Fig. 1A). Additionally, female mice had higher Tfhs in vLNs compared to male mice at day 14 and 21 post vaccination (Fig. 1A). To look at antigen experienced Tfhs, we gated on CD49d^+^CD11a^+^ T cells (gating strategy in Fig. S1A) (41, 42). Like total Tfhs, there was a significant increase in antigen experienced Tfhs in the vLNs of female but not male mice at 14 days post vaccination (Fig. 1B) and there were significant more of those antigen experienced Tfhs in female mice compared to male mice at that timepoint (Fig. 1B).

**Figure 1:**
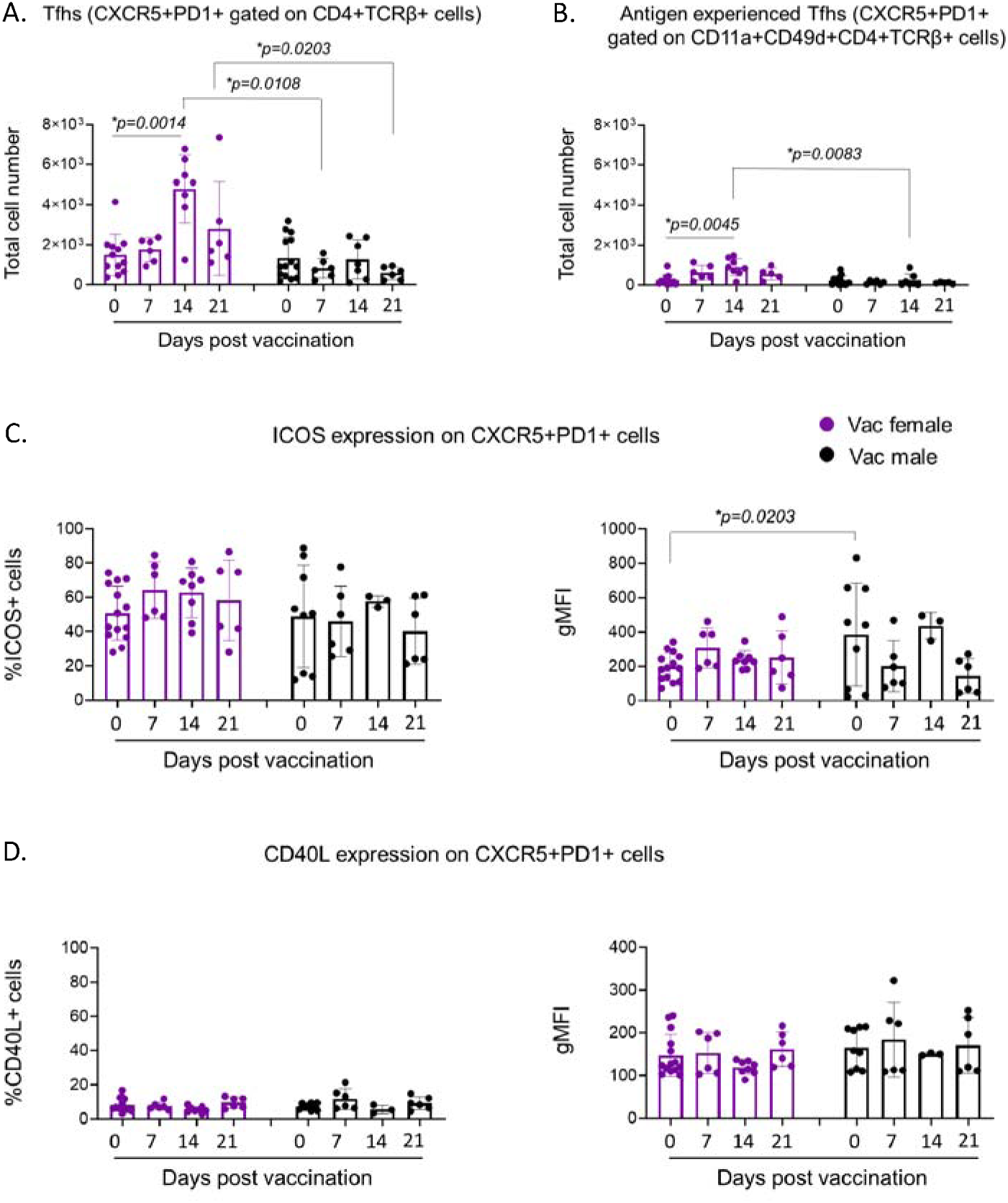
Sex-based differences in T follicular helper cells in the vaccine-draining lymph nodes following PCV vaccination. Adult C57BL/6 male and female mice were injected with PCV or PBS. Vaccine-draining lymph nodes were collected from all mice and assessed using flow cytometry. (A) Total and (B) antigen experienced T follicular helper cells (Tfh) were assessed at days 7, 14 and 21 post vaccination. Total Tfhs were assessed for their expression of (C) ICOS and (D) CD40L. (A) Total and (B) antigen experienced (CD11a^+^CD49d^+^) T cells (CD4^+^TCRβ^+^) were gated on and total numbers of CXCR5^+^PD1^+^ cells per mouse were determined. Total CXCR5^+^PD1^+^ cells were gated on and the percentage and geometric mean florescence intensity (gMFI) of (C) ICOS and (D) CD40L were determined. Data are pooled from 6 separate experiments and each dot represents an individual mouse. * denotes significant differences between the indicated groups as determined by (A&B) Kruskal-Wallis test followed by Dunn’s multiple comparisons test or (C) One-Way ANOVA followed by Šídák’s multiple comparisons test. Bar graphs represent the mean +/- SD.

An effective interaction between Tfhs and GC B cells requires multiple co-stimulatory molecules including Inducible T-cell costimulatory (ICOS) and CD40L (39, 40). We therefore investigated the expression of ICOS and CD40L on vLNs Tfhs on days 7, 14 and 21 following PCV vaccination (Fig. S1B). While there was no difference in the percentage expression of ICOS by Tfhs at all time points investigated, at baseline, Tfhs in female mice expressed more ICOS (gMFI) compared to male mice (Fig. 1C). No sex-based difference was observed in the percentage of CD40L expressing Tfhs or amount of CD40L expressed (Fig. 1D). No sex-based differences in total Tfhs, antigen experienced Tfhs or expression of ICOS and CD40L were observed in the spleen of male and female mice following vaccination (Fig.S2).

Next, we investigated B cell responses to PCV in male and female mice focusing particularly on germinal center B cells (GC) (B220^+^CD38^lo^CD95^+^) and plasmablasts (B220^+^CD138^+^) (Gating strategy in Fig. S3). Upon PCV vaccination, there was a gradual increase in the number of GC B cells in vLNs of female mice, with a difference from baseline at days 14- and 21-days post vaccination (Fig. 2A). However, such increase was not observed in vLNs of male mice (Fig. 2A). Similarly, an increase in plasmablasts was seen in vLNs of female mice with a significant peak at day 14 (Fig. 2B). Such increase was not observed in male mice and on day 21 post vaccination, female mice had more plasmablasts in vLNs than male mice (Fig. 2B). No sex-based differences in GC B cells or plasmablasts were observed in the spleen of male and female mice following vaccination (Fig. S4). Altogether, these data indicate that following PCV vaccination, female mice mount a robust Tfhs, GC B cells and plasmablast response in vLNs that is not observed in male mice.

**Figure 2:**
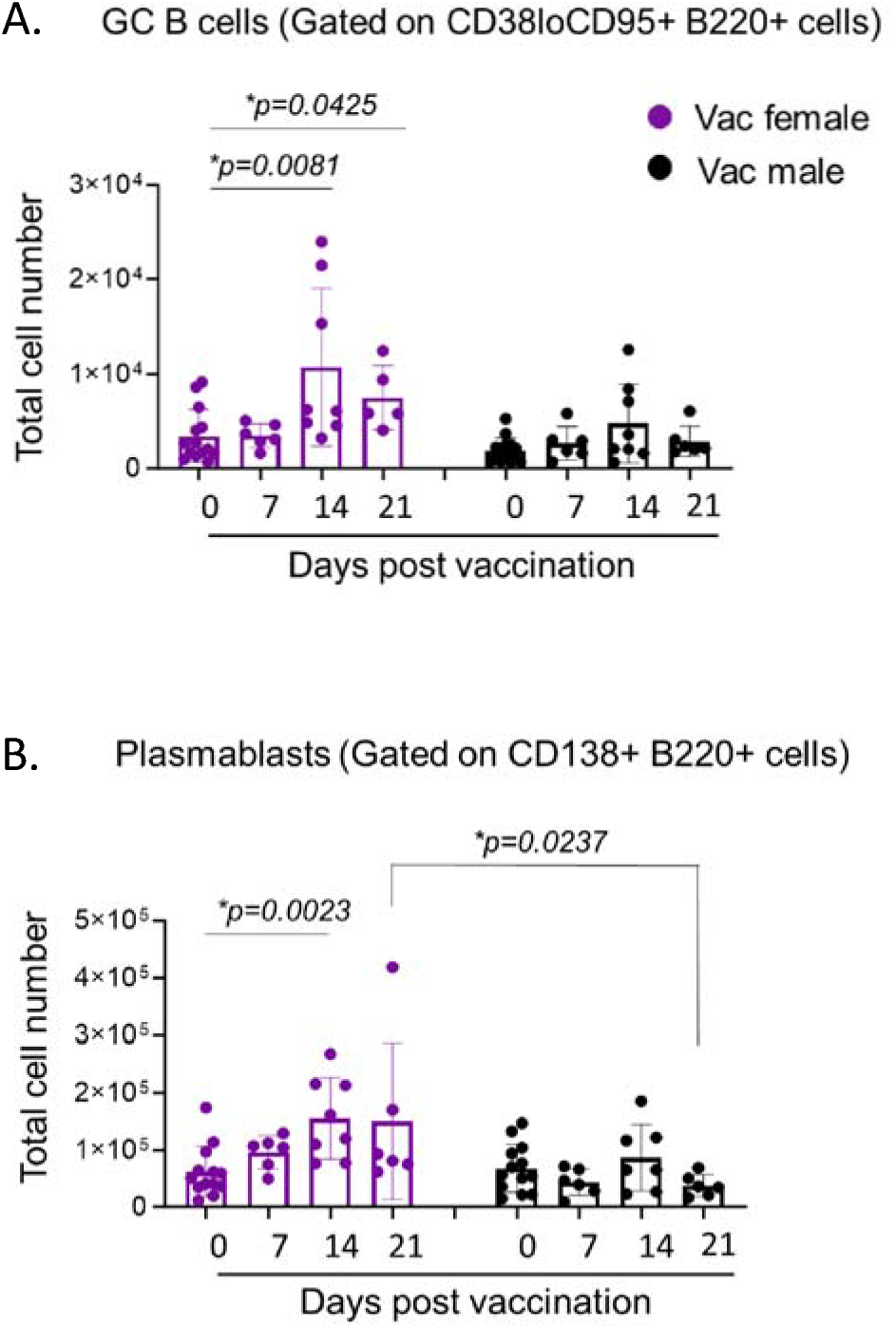
Sex-based differences in germinal center B cells and plasmablasts in the lymph nodes following PCV vaccination. Adult C57BL/6 male and female mice were injected with PCV or PBS. Vaccine-draining lymph nodes were collected from all mice and assessed using flow cytometry. (A) Germinal center (GC) B cells and (B) plasmablasts were assessed at days 7, 14 and 21 post vaccination. Total B220^+^ B cells were gated on and total numbers of (A) GC B cells (CD38^lo^CD95^+^) and (B) plasmablasts (CD138^+^) per mouse were determined. Data are pooled from 6 separate experiments and each dot represents an individual mouse. * denotes significant differences between the indicated groups as determined by Kruskal-Wallis test followed by Dunn’s multiple comparisons test. Bar graphs represent the mean +/- SD.

### Sex-based differences in circulating antibody responses following PCV vaccination

Next, we assessed the circulating antibody response in male and female mice following PCV vaccination. For that, sera collected prior to and on weeks 2 and 4 post vaccination were used to measure IgM and total IgG antibodies against heat-killed *S. pneumoniae* serotype 4 (TIGR4 strain) using an ELISA assay. Regardless of sex, mice mounted an IgM response to PCV post vaccination, with a significant increase from baseline at week 2 (Fig. 3A). However, male mice had significantly less antibodies during the peak IgM response at week 2 post vaccination compared to female mice (Fig. 3A). Similarly, mice mounted an IgG response post vaccination with isotype switching being observed by week 4 in both male and female mice. However, male mice had a significant 2-fold lower amounts of IgG at that timepoint compared to female mice (Fig. 3B).

**Figure 3:**
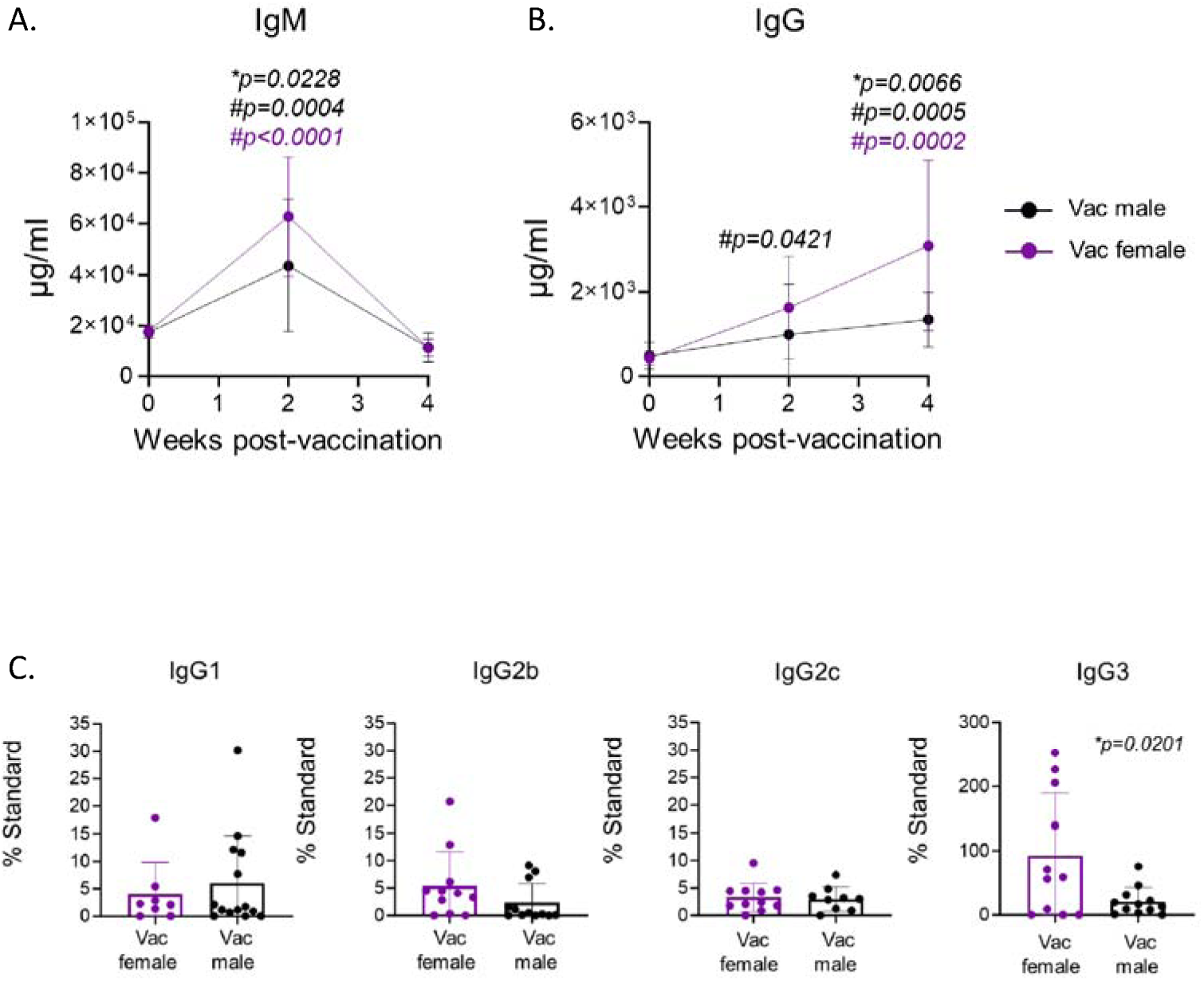
Sex-based differences in antibody responses following PCV vaccination. Adult C57BL/6 male and female mice were injected with PCV or PBS. At 0-, 2- and 4- weeks post vaccination, blood was collected from all mice and assessed for (A) IgM, (B) IgG and (C) IgG1, IgG2b, IgG2c and IgG3 antibodies against heat killed *S. pneumoniae* serotype 4 (TIGR4 strain) by ELISA. (A-B) Data are presented as concentrations in μg/ml and are pooled from a total of 3 experiments with at least 6 mice per group. * denotes significant differences compared to female mice at the same timepoint as determined by One-Way ANOVA followed by Šídák’s multiple comparisons and # denotes significant differences for the indicated group compared to their own baseline as determined by One-Way ANOVA followed by Dunnett’s multiple comparisons test. Line graphs represent the mean +/- 95% confidence interval. (C) Data are presented as percentage of standard sera and are pooled from a total of 3 experiments where each dot represents an individual mouse. * denotes significant differences between male and female mice as determined by Student’s t test. Bar graphs represent the mean +/- SD.

As there are different isotypes of IgG antibodies that can affect function (43), we assessed isotypes at week 4 post vaccination (the peak IgG response). Regardless of sex, mice mounted a much higher IgG3 antibody response compared to the other isotypes (Fig. 3C). However, we found that vaccinated male mice had significantly less IgG3 antibodies and slightly, but not significantly less IgG2b antibodies compared to vaccinated female mice (Fig. 3C). No differences were observed in levels of IgG1 and IgG2c antibodies (Fig. 3C). Together, these findings indicate that male mice make less antibodies compared to females in response to PCV and display changes in isotype switching.

### Sex-based differences in T cell subsets in the vaccine-draining lymph nodes following PCV vaccination

As isotype switching of antibodies by B cells is influenced by the cytokine environment (44) produced by different subsets of T cells (44, 45), we first investigated potential sex-based differences in T cell subsets following vaccination. On days 7, 14 and 21 post PCV vaccination, we assessed T cell responses in the spleen and vaccine draining lymph nodes (vLNs) of male and female mice. Overall, we found sex-based differences in the vLNs for all antigen-experienced T cells subsets (gating strategy in Fig. S5) investigated except for T helper 1 (Th1) cells (Fig. 4). When looking at antigen-experienced regulatory T cells (Tregs-FOXP3+CD11a+CD49d+ CD4^+^TCRβ^+^ T cells) (gating strategy in Fig. S5), we found that male mice had higher percentage at baseline but lower numbers at day 21 post vaccination compared to female mice (Fig. 4A). For antigen-experienced Th17 (RORγT+FOXP3-CD11a+CD49d+ CD4^+^TCRβ^+^T cells) (gating strategy in Fig. S5) cells, male mice had lower numbers on days 7 and 21 post vaccination compared to female mice with no difference observed in the respective percentages (Fig. 4B). In contrast to Tregs and Th17, male mice had higher numbers of antigen-experienced Th2 (GATA3+FOXP3-CD11a+CD49d+ CD4^+^TCRβ^+^ T cells) (gating strategy in Fig. S5) cells at day 21 post vaccination compared to female mice with no difference observed in the percentages (Fig. 4C). No sex-based differences were observed in both the percentage and numbers of Th1 (Tbet+FOXP3-CD11a+CD49d+ CD4^+^TCRβ^+^ T cells) (gating strategy in Fig. S5) cells (Fig. 4D). No sex-based differences in any of the T cell subsets investigated were observed in the spleen of male and female mice following vaccination (Fig. S6). However, when we assessed the composition of the cytokine milieu in the spleen, we found significant changes between vaccinated male and female mice (Fig. S7). For that, spleens obtained on days 7 and 14 post vaccination were used to assess pan-cytokine/chemokine levels using multiplex immunoblots. Comparing the fold change of responses observed in males over females, we found that for most cytokines tested, male mice showed similar responses compared to female mice at day 7 post vaccination (Fig. S7). Exceptions included cytokines that affect B cell responses (44) including IL-33, TNF-α, IL-15, IL-7 and the IL-6 family member leukemia inhibitory factor (LIF), that were 5-10-fold lower in male mice compared to female mice. In contrast, IL-10, CRP and CCL19 were 10-fold higher while CCL6 and C5a were more than 100-fold elevated in male mice. However, by day 14, male mice had exacerbated responses in almost all factors measured (Fig. S7) including more than 100-fold higher levels of a wide range of chemokines, Th1 cytokines including IFNγ, Th2 cytokines including IL-10, IL-13 and IL-4 and Th17 related cytokines such IL-22 (Th17-induced) and IL-23 (Th17 inducing) (45). Male mice also had higher levels of factors that activate B cells including B cell activating factor (BAFF) (46) and Pentraxin 3 (PTX3) (47) (10-fold higher) as well as IL-5 (100-fold higher) (48, 49). These data suggest that there are sex-based differences in T cell responses and the cytokine environment following PCV vaccination that can influence the antibody response.

**Figure 4:**
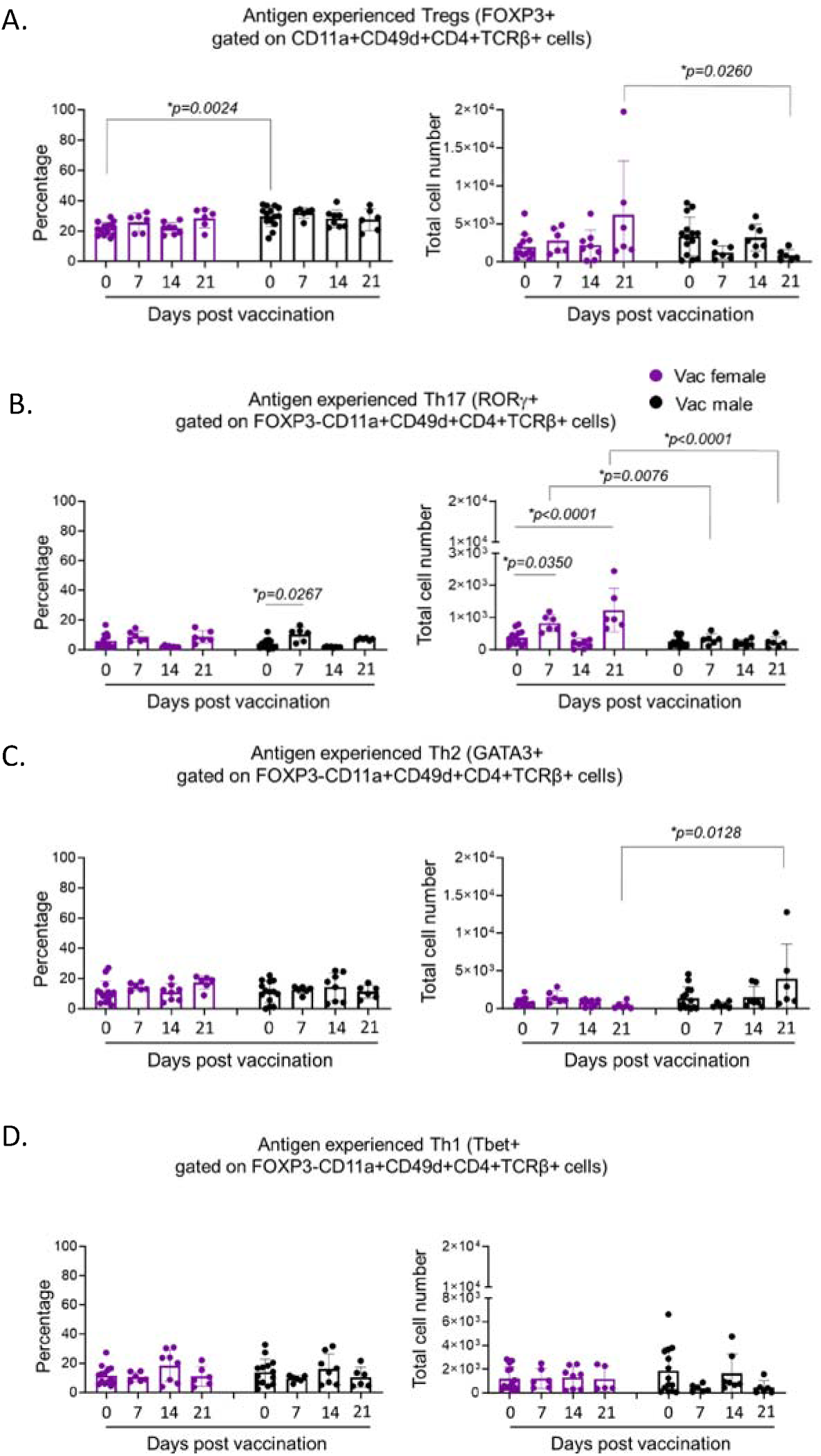
Sex-based differences in T cell subsets in the lymph nodes following PCV vaccination. Adult C57BL/6 male and female mice were injected with PCV or PBS. Vaccine-draining lymph nodes were collected from all mice and assessed using flow cytometry. (A-D) Antigen experienced T cells (CD11a^+^CD49d^+^CD4^+^TCRβ^+^) were gated on and percentages (left) and numbers (right) of T cell subsets per mouse were determined at days 7, 14 and 21 post vaccination. (A) Regulatory T cells (FOXP3^+^), (B) T helper 17 (RORγT^+^FOXP3^-^), (C) Th2 (GATA3^+^FOXP3^-^) and (D) Th1 (Tbet^+^FOXP3^-^) were gated on. Data are pooled from 6 separate experiments and each dot represents an individual mouse. * denotes significant differences between the indicated groups as determined by (A&C) Kruskal-Wallis test followed by Dunn’s multiple comparisons test or (B) One-Way ANOVA followed by Šídák’s or Dunnett’s multiple comparisons test. Bar graphs represent the mean +/- SD.

### S. pneumoniae are bound with lower amounts of antibodies upon exposure to sera from vaccinated males

As antibody function is crucial for vaccine efficacy (50), we wanted to assess the antimicrobial function of the sera in PCV vaccinated male and female mice. For that, sera collected at 4 weeks post PCV vaccination from male and female mice was used to perform different functional assays. Since there was approximately a 2-fold difference in the amount of IgG present in the blood of vaccinated male versus female mice, we used different sera percentages to compare across sexes. First, we investigated the ability of antibodies in the sera to bind the surface of *S. pneumoniae*. GFP-expressing *S. pneumoniae* serotype 4 were incubated with sera at 1-, 3- or 6% of total assay volume, and the degree of binding was detected using flow cytometry (Fig. S8). We found that within a given sera dilution, regardless of sex, there was an upregulation in antibody binding to bacteria when using vaccinated sera compared to naïve sera (Fig. 5). Additionally, within each sera percentage, less antibodies bound to bacteria upon incubation with vaccinated male sera compared to vaccinated female sera, with a significant difference in the 3% condition (Fig. 5). However, when the amounts of antibody present in the sera of male and female mice are matched, (comparing 3% vaccinated female sera with 6% vaccinated male sera, a 2-fold difference), there was no difference in binding of antibodies to bacteria (Fig. 5 & Table S1) across the sexes. These data show that the lower antibody levels observed in vaccinated male mice (Fig. 3B) result in less antibodies being bound to *S. pneumoniae* upon sera exposure.

**Figure 5:**
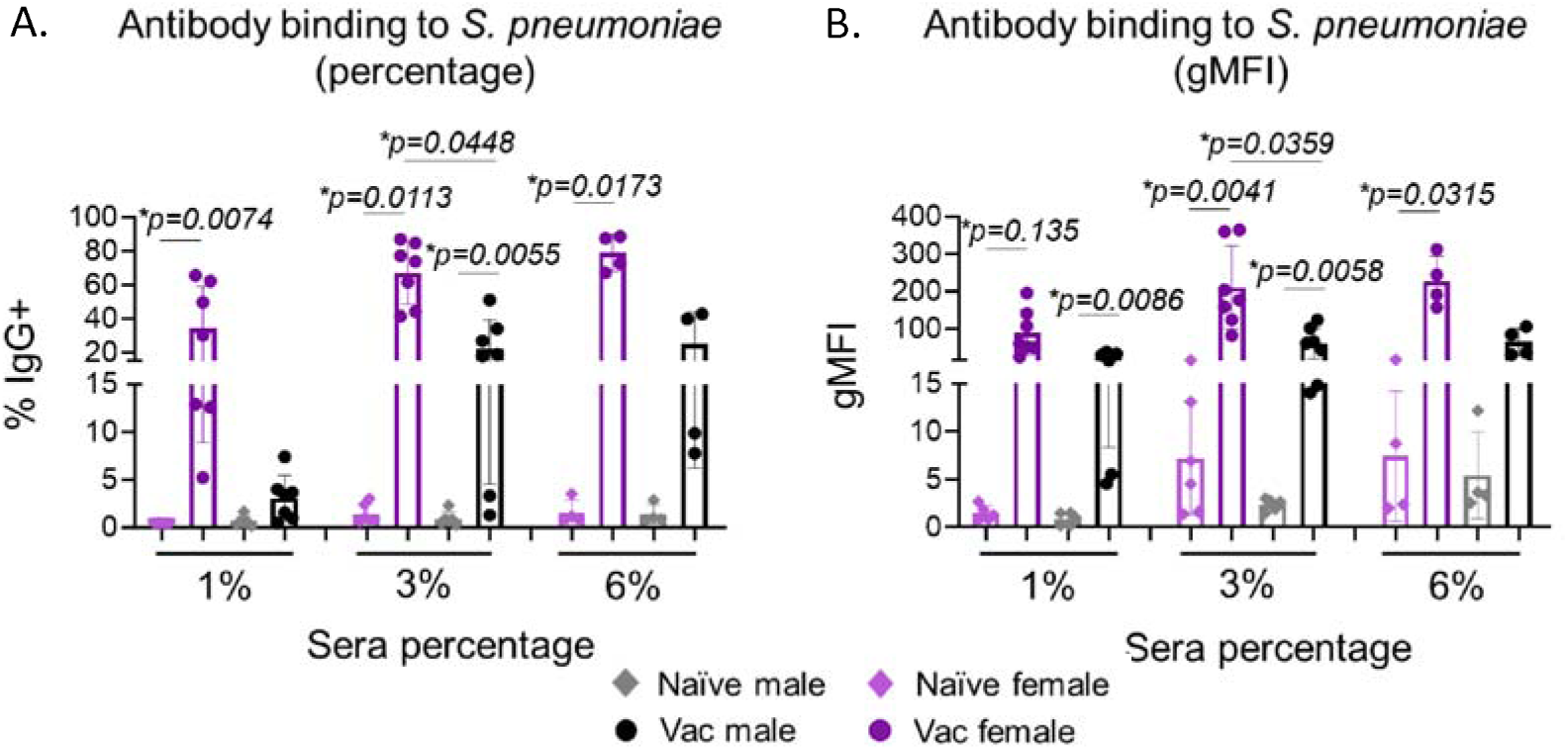
Less antibodies bind to *S. pneumoniae* upon incubation with sera from vaccinated males. Adult C57BL/6 male and female mice were injected with PCV or PBS. Sera collected 4-weeks post vaccination were incubated with the *S. pneumoniae* at 1-, 3- and 6% dilutions to assess the binding of antibodies to bacterial surfaces via flow cytometry. *S. pneumoniae* were incubated with the indicated sera conditions, washed, and labeled with fluorescently tagged anti-mouse IgG. (A) Percent and (B) gMFI of IgG bound to *S. pneumoniae* are shown. Data are pooled from 7 separate experiments with each dot representing the average of technical replicates from each experiment. * denotes significant differences between the indicated groups as determined by Kruskal-Wallis test followed by Dunn’s multiple comparisons test. Bar graphs represent the mean +/- SD.

### Sera from vaccinated males are less efficient at neutralizing binding of *S. pneumoniae* to pulmonary epithelial cells

One of the ways antibodies protect the host is by neutralization of bacterial binding to epithelial cells, which is required for establishing pulmonary infection (34, 51–53). To examine neutralization, we used an established *in vitro* binding assay using the human lung epithelial cell line H292, a mix of type I and II pneumocytes (32). We compared the ability of sera from vaccinated male and female mice to block bacterial binding to H292 cells relative to naïve controls for each sex. Due to differences in antibody levels in the sera, we tested varying concentrations. We found that at 10%, sera from vaccinated male mice did not prevent binding of *S. pneumoniae* to H292 cells as opposed to a significant, 37% reduction in binding observed with sera from vaccinated female mice (Fig. 6). Additionally, this reduction in bacterial binding was higher in female mice compared to male mice (Fig. 6). At 20% dilution, while both sera from vaccinated male and female mice lead to a reduction in bacterial binding, sera from vaccinated females neutralized binding better than sera from vaccinated males (Fig. 6). Lastly, when comparing 10% vaccinated female sera to 20% vaccinated male sera (to match the amount of antibodies), the efficiencies in *S. pneumoniae* neutralization were similar (Fig. 6 and Table S1). This suggests that the observed sex-based differences in neutralization capacity of the sera could be attributed to the difference in amount of antibodies present in the sera post PCV vaccination (Fig. 3B).

**Figure 6.**
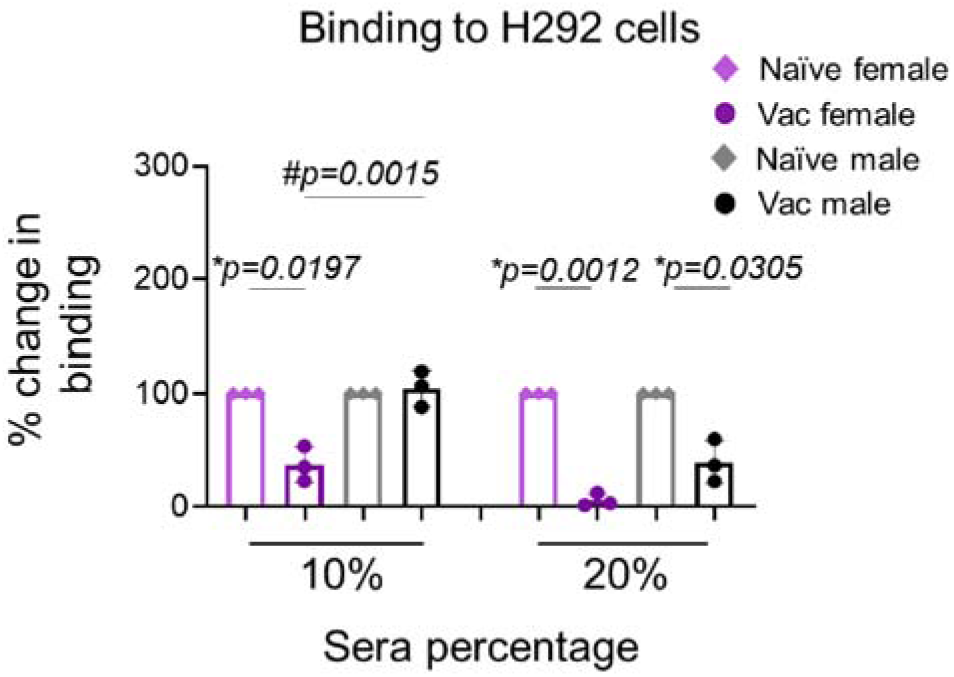
Sera from vaccinated males are less efficient at neutralizing binding of *S. pneumoniae* to H292 cells. Adult C57BL/6 male and female mice were injected with PCV or PBS. Sera were collected at 4-weeks post vaccination and assessed for the ability to neutralize bacterial binding to mammalian cells. *S. pneumoniae* were pre-opsonized with sera at 10- and 20% dilutions for 45 minutes and then incubated with H292 cells for 1 hour at an MOI of 10. The number of bound bacteria was measured by plating on blood agar plates and the percent bacterial binding calculated with respect to bacteria incubated with sera alone for each opsonization condition. The effect of sera on bacterial binding was then determined relative to the naive group for each sex. Data are pooled from 3 separate experiments with each dot representing the average of technical replicates from each experiment. * denotes significant differences from the naive groups as determined by One sample t test and # denotes significant differences between the indicated groups as determined One-way ANOVA followed by Šídák’s multiple comparisons test. Bar graphs represent the mean +/- SD.

### No sex-based differences in the ability of sera from vaccinated hosts to promote opsonophagocytic uptake and killing of *S. pneumoniae* by neutrophils

The opsonic capacity of antibodies is a key correlate of vaccine efficacy (54). Antibodies allow efficient bacterial uptake and phagocytosis by neutrophils (also known as polymorphonuclear cells or PMNs), which is key for protection of vaccinated hosts (55). We first investigated the ability of sera from vaccinated hosts to mediate opsonophagocytic uptake of *S. pneumoniae* by PMNs using GFP-expressing *S. pneumoniae* and flow cytometry and inside-out staining (36). We found that overall, there was no difference in the percent of bacteria associated with PMNs (total of intracellular and extracellular bacteria) upon incubation with bacteria opsonized with sera from naïve or vaccinated male and female mice at any of the dilutions tested (Fig. S9A). Similarly, there was no difference in the percent of intracellular bacteria between the different groups evaluated (Fig. S9B).

We then looked at the ability of sera to induce opsonophagocytic killing of *S. pneumoniae* by PMNs. There have been reports of sex-based differences in PMN phagocytosis (56, 57), therefore, to assess only the functionality of the sera, we kept the PMNs constant. PMNs from naïve female mice were infected with *S. pneumoniae* pre-opsonized with different concentrations of sera from naïve or vaccinated from male and female mice. We found that there was an increase in opsonophagocytic killing when using sera from vaccinated mice compared to naïve controls at 3% for both females and males and no differences across the sexes (Fig. 7A). When comparing matched antibody amounts conditions (3% vaccinated female sera vs 6% vaccinated male sera), there was no difference in level of opsonophagocytic killing in females compared to males (Fig. 7A & Table S1).

**Figure 7.**
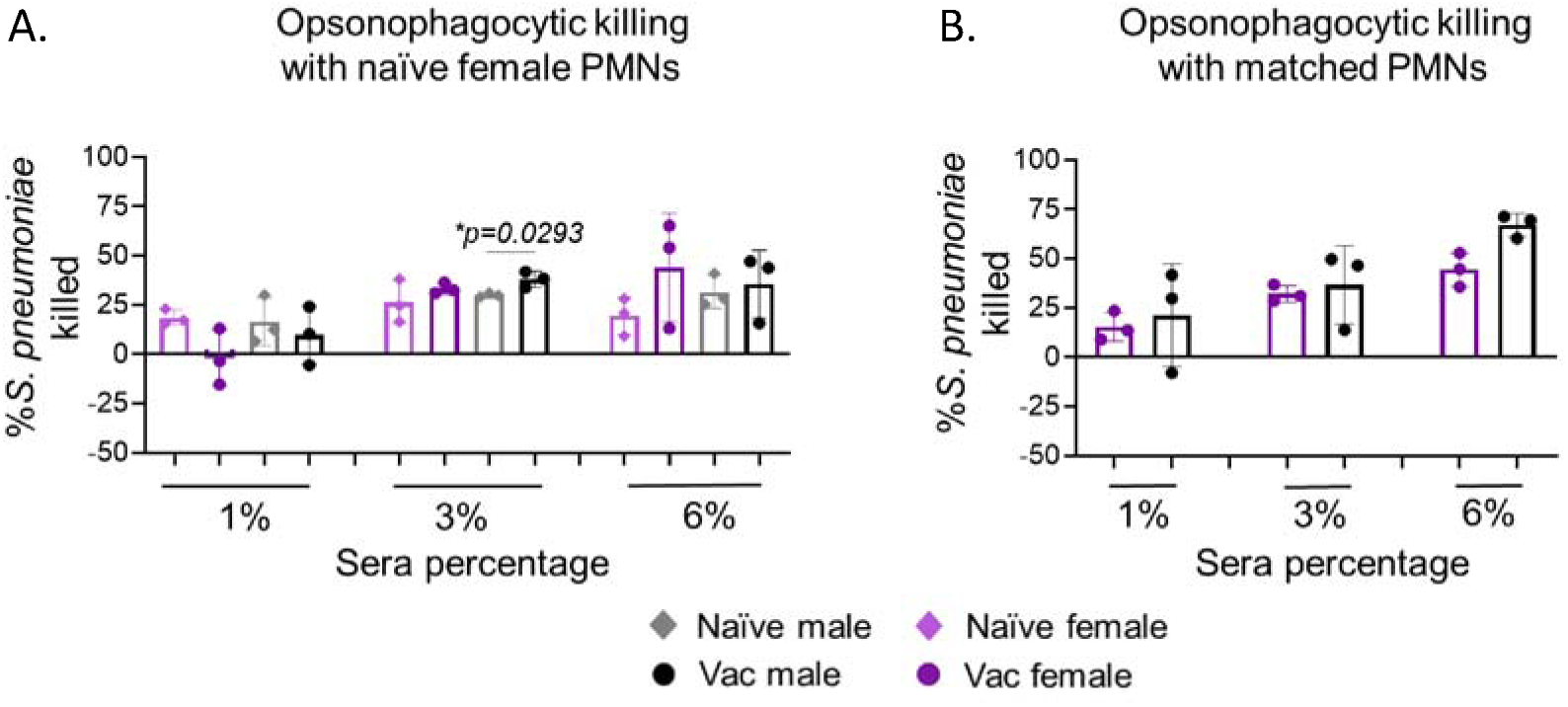
No sex-based differences in the ability of sera from vaccinated mice to mediate opsonophagocytic killing of *S. pneumoniae* by PMNs. Adult C57BL/6 male and female mice were injected with PCV or PBS. Sera collected 4-weeks post vaccination in all mice were assessed for their ability to induce opsonophagocytic killing by PMNs. *S. pneumoniae* were opsonized with sera at 1-, 3- and 6% dilutions and incubated with (A) PMNs from naïve females or (B) matched with PMNs from the same group. Percent bacterial killing was determined with respect to a no PMN control for each sera condition. Data are pooled from 3 separate experiments with each dot representing the average of technical replicates from an individual mouse. * denotes significant differences between the indicated groups as determined by (A) student t test or (B) One-way ANOVA followed by Šídák’s multiple comparisons test. Bar graphs represent the mean +/- SD.

To assess a more physiologically relevant condition, PMNs from vaccinated male and vaccinated female mice were incubated with *S. pneumoniae* pre-opsonized with matched sera from the same host. We observed a dose dependent increase in bacterial killing in both male and female mice across the sera dilutions (Fig. 7B). However, there was no difference in bacterial killing between males and females at any of the sera dilutions tested (Fig. 7B). The above data suggest that there is slight, but not significant sex-based differences in the opsonic capacity of antibodies in the sera of PCV vaccinated hosts.

### Sera from vaccinated males are less efficient at direct killing of *S. pneumoniae*

Antibody binding to bacteria also triggers complement deposition on the bacterial surface, leading to complement mediated bacterial lysis (58, 59). To test complement deposition on the surface of *S. pneumoniae* in the presence of sera from vaccinated hosts, we used a flow cytometry-based assay. *S. pneumoniae* were incubated with varying concentrations of sera and the degree of C3 binding (a key component of the complement system (59)) detected using flow cytometry (gating strategy in Fig. S10). We observed a dose dependent increase in C3 deposition on *S. pneumoniae* across 1-, 3- and 6% sera dilutions for both male and female mice (Fig. 8A). However, there was no difference in C3 deposition between sera from naïve and vaccinated or male and female mice (Fig. 8A). This was not surprising given that IgM, which is present in naïve sera, is significantly more efficient at complement fixation compared to IgG (60) and could mask any vaccine-mediated effects.

**Figure 8.**
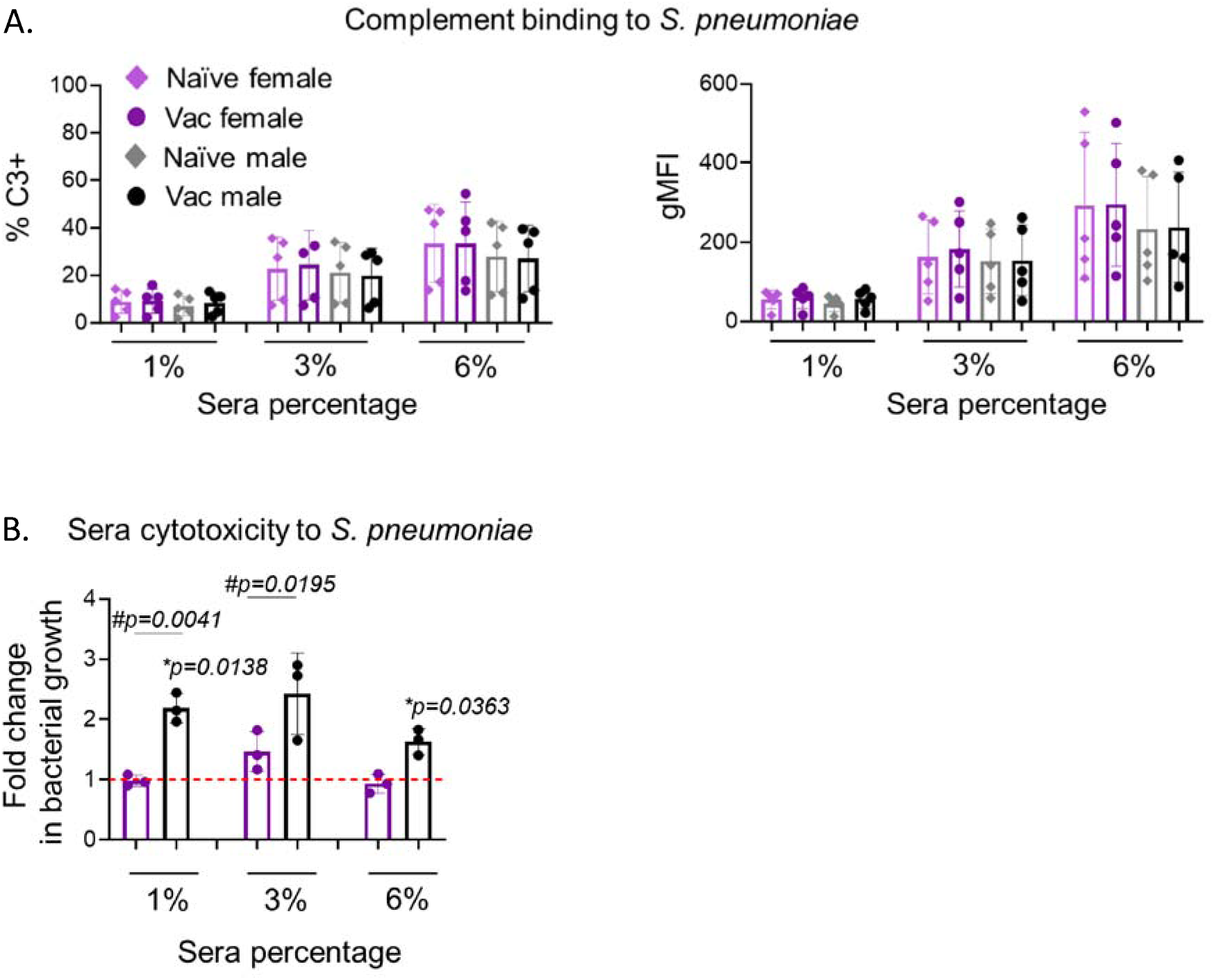
Sera from vaccinated males are less efficient at direct killing of *S. pneumoniae*. Adult C57BL/6 male and female mice were injected with PCV or PBS. Sera collected 4-weeks post vaccination were incubated with the pneumococcus at 1-, 3- and 6% dilutions to assess their ability to (A) mediate complement deposition on the bacterial surface via flow cytometry and (B) induce bacterial cytotoxicity. (A) *S. pneumoniae* were incubated with the indicated sera, washed, and labeled with fluorescently tagged anti-C3 antibody. Percent and gMFI of C3 bound to *S. pneumoniae* are shown. (B) *S. pneumoniae* were incubated with the indicated sera and plated on agar plates. Fold change in bacterial viability from input is shown where values above and below 1 indicate increased or decreased bacterial net growth respectively.. (A) Data are pooled from 5 separate experiments with each dot representing the average of technical replicates from each experiment. (B) Data are pooled from 3 separate experiments with each dot representing the average of technical replicates from each experiment. * denotes significant differences of the indicated groups from 1 as determined by One Sample t test and # denotes significant differences between the indicated groups as determined by One-way ANOVA followed by Šídák’s multiple comparisons test. Bar graphs represent the mean +/- SD.

We then looked at sera-mediated cytotoxicity by determining the number of bacteria that survived a 40-minute incubation with different dilutions of sera from vaccinated hosts. We found that within each sera dilution, sera from vaccinated males could not control the growth of *S. pneumoniae* compared to sera from vaccinated females and bacterial growth was observed in the presence of sera from male mice (Fig. 8B). Additionally, at 3% sera dilution, there was a significant difference in control of bacterial growth between males and females (Fig. 8B). This shows that sera from vaccinated males are less cytotoxic to *S. pneumoniae* compared to female counterparts. Lastly, when comparing 3% sera from vaccinated females to 6% sera from vaccinated males (matched amount of antibodies), the efficiencies in controlling *S. pneumoniae* growth were similar (Fig. 8B). This suggests that the difference in the ability of the sera to control bacterial growth across the sexes is due to the overall lower antibody amount in the sera of vaccinated male mice (Fig. 3B).

### PCV vaccinated male mice are not as protected against invasive pneumococcal infection as vaccinated females

With the differences observed in antibody response and function, we next evaluated the level of protection offered by PCV in male and female mice *in vivo*. Four weeks following PCV vaccination, male and female mice were infected intratracheally with 10^7^ colony forming unit (CFU) of the invasive *S. pneumoniae* TIGR4 strain (vaccine-covered serotype 4). Mice were either monitored for disease progression over the course of 7 days or assessed for bacterial burden in their lungs and blood 24 hours post infection. When looking at survival post infection, regardless of sex, all naïve mice succumbed to the infection within the first 48 hours (Fig. 9A). Vaccinated female mice were significantly protected compared to naïve controls. Enhancement of protection was also observed following vaccination of male mice, although this was not significant (Fig. 9A). Further, while the majority (85%) of vaccinated female mice survived, only half of vaccinated male mice survived the infection, with female mice displaying significantly better survival compared to male mice (Fig. 9A). This difference in protection was reflected in their clinical scores at 24 hours post infection, which are a set of criteria that determine disease severity and include assessment of weight loss, posture, activity and respiration (37). While a small percentage of vaccinated female mice had a score higher than 0, nearly all vaccinated male mice were sick, with >75% of them displaying signs of disease (Fig. 9B). Looking at bacterial burden 24 hours post infection, vaccinated female mice were able to control bacterial numbers in their lungs (Fig. 9C) as well as spread to the blood as opposed to male mice which showed no improvement in pulmonary bacterial control and only 50% protection against systemic spread upon vaccination (Fig. 9D). These data demonstrate that male mice remain more susceptible to invasive pneumococcal infection even following vaccination with PCV and are less protected than females.

**Figure 9.**
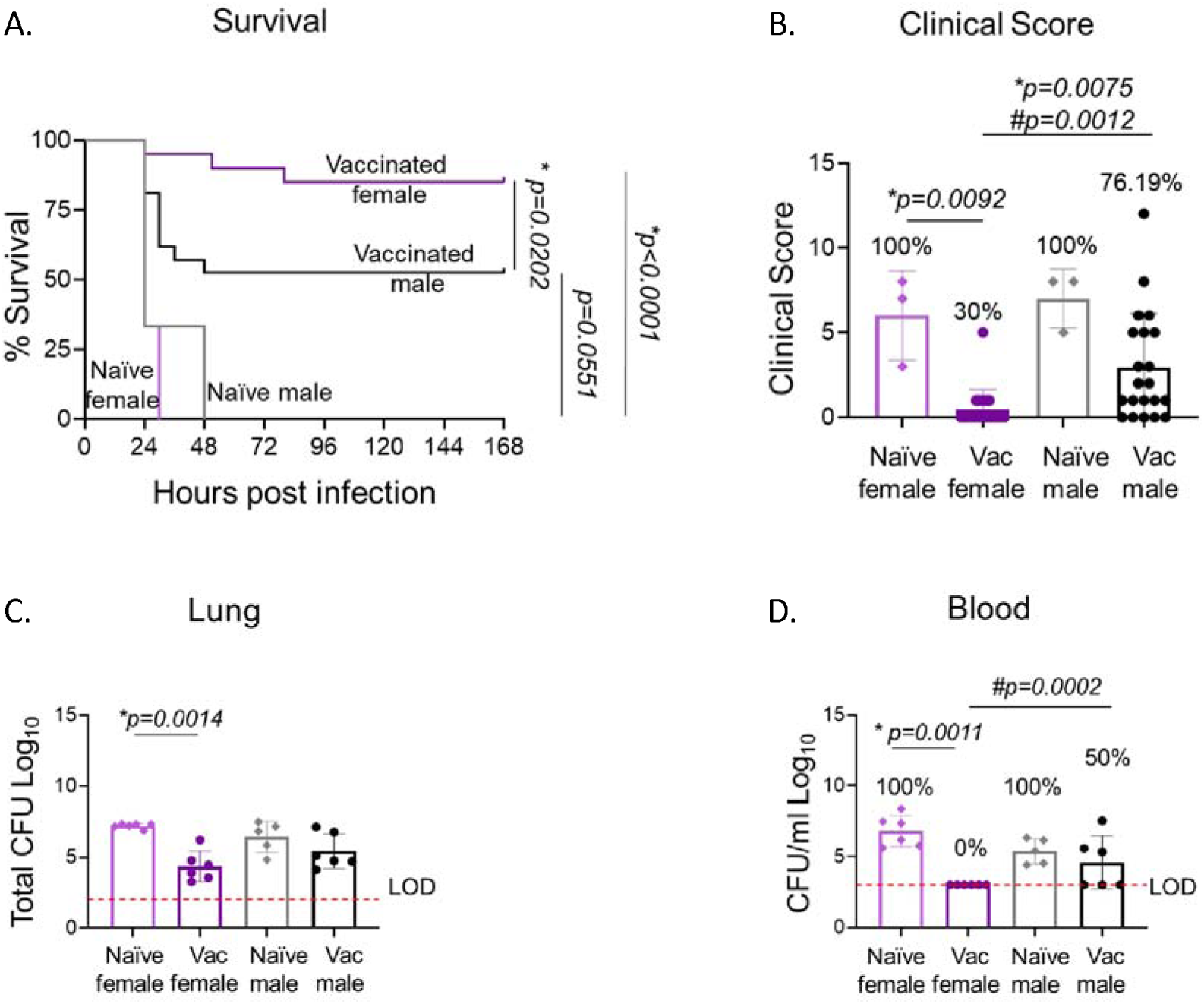
PCV vaccinated male mice are not as protected from invasive pneumococcal infection as vaccinated females. Adult C57BL/6 male and female mice were injected with PCV or PBS and infected 4 weeks post vaccination with 10^7^ CFU of (A-D) *S. pneumoniae* TIGR4 (serotype 4). For one set of experiments, (A) survival of the mice was monitored over 7 days post infection and (B) disease severity from those mice was assessed 24 hours post infection. For a second set of experiments, bacterial burden in the (C) lung and (D) blood of the mice was assessed 24 hours post infection. (A-B) Data are pooled from 6 separate experiments with a total of n=3 naïve females, n=3 naïve males, n=21 vaccinated females and n= 21 vaccinated males and (B) each dot represents an individual mouse. * denotes significant differences between the indicated groups as determined by (A) Mantel-Cox or (B) Kruskal-Wallis test followed by Dunn’s multiple comparisons test. # denotes significant differences in disease incidence between the indicated groups as determined by Fisher’s exact test. (C-D) Data are pooled from separate experiments and each dot represents an individual mouse. * denotes significant differences between the indicated groups as determined by Kruskal-Wallis test followed by Dunn’s multiple comparisons test. # denotes significant differences in bacteremia incidence between the indicated groups as determined by Fisher’s exact test. Bar graphs represent the mean +/- SD.

### PCV vaccinated male mice are not as protected from lung-localized pneumococcal infection as vaccinated females

Finally, we wanted to assess protection of PCV vaccinated male and female mice against a lung-localized strain of *S. pneumoniae.* To test that, we measured protection against infection with *S. pneumoniae* strain EF3030, a vaccine-covered serotype 19F otitis media strain that does not disseminate to the blood in murine models (61). Vaccinated male and female mice were intratracheally infected 4 weeks post PCV vaccination with EF3030 and bacterial burden was assessed 30 hours later. As seen with invasive infection, vaccinated female mice were able to control bacterial numbers in the lung and showed a significant, 100-fold lower burden compared to naïve controls (Fig. 10). However, no difference in bacterial burden was observed in the lung of male mice following vaccination (Fig. 10). Consistent with this being a lung-localized infection, no bacteria were detected in the blood of any of the mice. These data demonstrate that PCV fails to protect male mice against lung localized pneumococcal infection as efficiently as female mice.

**Figure 10.**
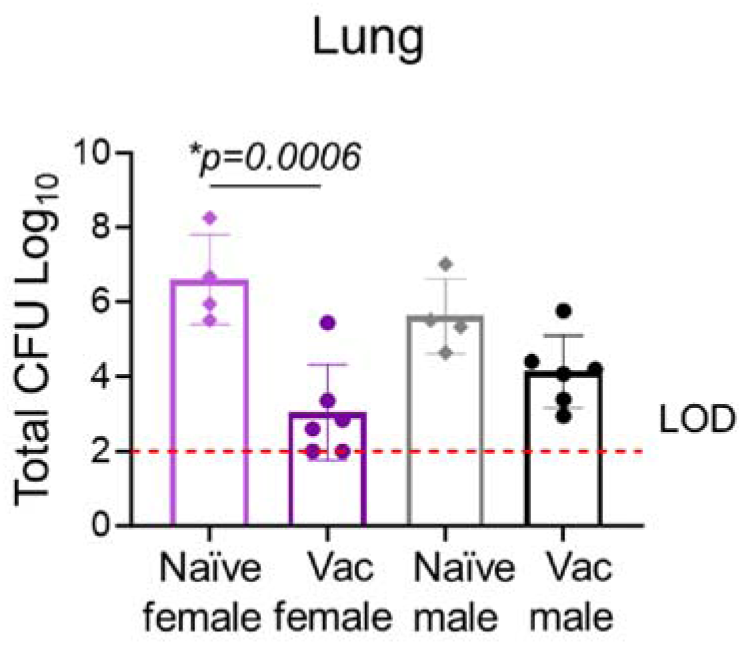
PCV vaccinated male mice are not as protected from a lung-localized infection as vaccinated females. Adult C57BL/6 male and female mice were injected with PCV or PBS. At 4 weeks post vaccination, mice were infected with 10^7^ CFU of *S. pneumoniae* EF3030 (serotype 19F) and bacterial burden in the lung were assessed 24 hours post vaccination. Each dot represents an individual mouse. * denotes significant differences between the indicated groups as determined by Kruskal-Wallis test followed by Šídák’s multiple comparisons test.

## Discussion

Sex-based differences in humoral responses have been widely reported with studies showing that adult males produce lower levels of antibodies to several vaccines (62, 63) including those against pulmonary pathogens such as *S. pneumoniae* (26), SARS-CoV-2 (64) and influenza (5). Sex-based factors refer to biological determinants that include sex chromosomes, reproductive organs and sex hormones that have all been reported to influence immune responses (65, 66). In this study, we found that there are sex-based differences in both humoral and cellular immune responses to the pneumococcal conjugate vaccine with reduced numbers of several cell types involved in antibody production, namely T follicular helper cells, germinal center B cells and plasmablasts in adult male mice compared to female mice. This was associated with reduced antibody levels and reduced vaccine efficacy against pneumococcal infection in adult male versus female mice. The reduction in antibody production (26, 67, 68) and vaccine-mediated protection (24, 69–71) in males recapitulates what has been reported in humans, and can be used to dissect the sex-based variables driving these differences.

Surveillance studies have reported that in the USA, the incidence of pneumococcal disease remans higher in males compared to females even after the introduction of vaccines (24, 69–71). The efficacy of the 23-valent pneumococcal polysaccharide vaccine, licensed for use in older adults since the 1980s, differs between the sexes where overall reduction of mortality and protection against pneumonia is higher in females above the age of 65 years, compared to males (24, 69–71). The pneumococcal conjugate vaccine first introduced in 2000 and recommended for children and older adults as well as those at risk was similarly associated with reduced all-cause mortality in females 65-69 years old but not in males (71). PCV was also associated with lower incidence of IPD in adult females and with lower incidence of IPD in girls compared boys across several studies (24, 69–71). Here, we found that PCV was better able to protect adult female mice against lung-localized infection as well as systemic spread compared to male mice. As differences in PCV efficacy is observed in infants and young children prior to puberty, it is likely that sex chromosomes play an important role in vaccine-mediated protection. The X-chromosome encodes several genes that control immunity including several TLRs, regulatory T cell transcription factor FOXP3, B and T cell costimulatory molecule CD40L, among others (72). As females express two copies of the X chromosome (XX) while males express one (XY), gene composition as well as incomplete inactivation of X-linked immune genes (65) may play a role in differences in PCV efficacy observed across the sexes. A study tracking the effect of sex chromosomes vs sex hormones on immune responses against immunization with an acapsular heat-killed *S. pneumoniae* found differential effects on humoral responses (73). Using the Four Core Genotype mice, that allow formation of ovary bearing female mice with XX or XY chromosomes as well as testes-bearing male mice with XX or XY chromosomes, demonstrated that while female gonadal sex enhanced production of antigen specific B cells, the XX chromosome enhanced the number of CD138+ plasma cells independent of gonadal sex or sex hormones (73). Other genetic factors likely have important roles in PCV responses as well, as some populations in the Netherlands do not display the sex-based differences in PCV-mediated protection reported in the USA (24).

In tracking antibody response, we found that male mice produce significantly less anti-pneumococcal antibodies in response to PCV compared to female mice. This was observed in both IgM levels, as well as IgG. Sex-based differences in anti-pneumococcal antibodies are well established and have been reported across several studies (26, 28, 30, 67, 68). In non-vaccinated adults (ages 18-65), naturally acquired IgM against polysaccharides across several pneumococcal serotypes, including serotype 4 that we focused on in this study, were reported to be higher in females compared to males (74). Similarly, naturally acquired IgG against a few pneumococcal serotypes are higher in adult human females compared to males (74, 75). For pneumococcal vaccines, most studies in adults have been done with PPSV, with opposing reports showing sex-based differences in IgG levels that were higher either in males or females (26, 28, 30). Fewer studies have examined IgG responses to the pneumococcal conjugate vaccine in adults and most studies focusing on children have shown significantly lower levels of IgG against several bacterial capsular serotypes in males compared to females, including for serotypes 4 and 19F (67). This is consistent with the lower amounts of anti-serotype 4 polysaccharides antibodies and the lower protection against infections with serotype 4 and 19F strains that we reported here. When assessing functionality, we found that sera from vaccinated female mice had better opsonic, cytotoxic, and bacterial neutralizing capacity. This was due to the higher amounts of antibodies in the sera and is consistent with improved opsonic capacity reported for sera from vaccinated girls compared to boys (67). We further found changes in isotype switching to IgG3 in females compared to males and mouse IgG3 (equivalent to human IgG2) is known to respond to bacterial polysaccharides (76, 77), and protect mice against pneumococcal infection (78).

In exploring mechanisms, we found sex-based differences in cell types involved in antibody production, with higher number T follicular helper cells, germinal center B cells and plasmablasts in female mice compared to male mice. Prior studies have similarly found increase in Tfh cell number in response to immunization with T-cell dependent antigens in females compared to males (79, 80). In humans, PCV vaccination elicited peripheral Tfh cells that correlated with increase in production of opsonic antibodies against pneumococci (81), suggesting that Tfhs are important for B cell function in response to the pneumococcal conjugate vaccine. Higher number of germinal center B cells in response to influenza vaccination have also been reported in females compared to males (82, 83) and estrogen was shown to directly elicit production of activation-induced deaminase (AID), the enzyme required for somatic hypermutation and class switch recombination in B cells (84). Estrogen was shown to induce class switching to IgG3 (84), which is in line with higher production of IgG3 in female mice we observed here. Sex-based differences in other B cell subsets including higher levels of antibody producing cells (as observed here) and innate B-1 cells in females compared to males have also been reported (73, 82, 85, 86). We also found changes in T cell subsets in response to PCV, with lower number of Tregs and Th17, but higher number of Th2 cells in the vaccine-draining lymph nodes of male mice compared to female mice. We further found changes in the splenic cytokine environment. Prior studies have demonstrated clear sex-based differences in T cell polarization and cytokine production in several infection, allergic, and autoimmune models that were driven by differences in sex hormone levels (87–93). Production of regulatory T cells was also shown to be enhanced by estrogen, and during pneumococcal pneumonia, female mice were reported to have higher Treg numbers in the lungs compared to male mice (94, 95). These sex-based changes in the T and B cell compartments are likely controlling the lower Ab production and changes in isotype switching we observed in male versus female mice.

In summary this study demonstrates sex-based differences in immune responses to PCV and can be used as a tool for testing improved vaccine strategies in the future. One caveat is the use of adult mice since PCV is recommended in people at the extremes of age i.e., children and older adults or adults with underlying conditions (31). However, this model can be used for improving vaccine design and dissecting the contribution of sex chromosomes vs sex hormones to pneumococcal vaccine responses and efficacy against infections moving forward. Future studies will explore the intersection of sex and age in pneumococcal vaccine responses.

## Author Contributions

EYIT and AB conducted research, analyzed data, and wrote the paper. AYK conducted research and analyzed data. MB conducted research. ENBG designed research, wrote the paper, and had responsibility for final content. All authors read and approved the final manuscript.

