## Supplemental Data for "Sex-based difference in immune responses and efficacy of the pneumococcal conjugate vaccine"

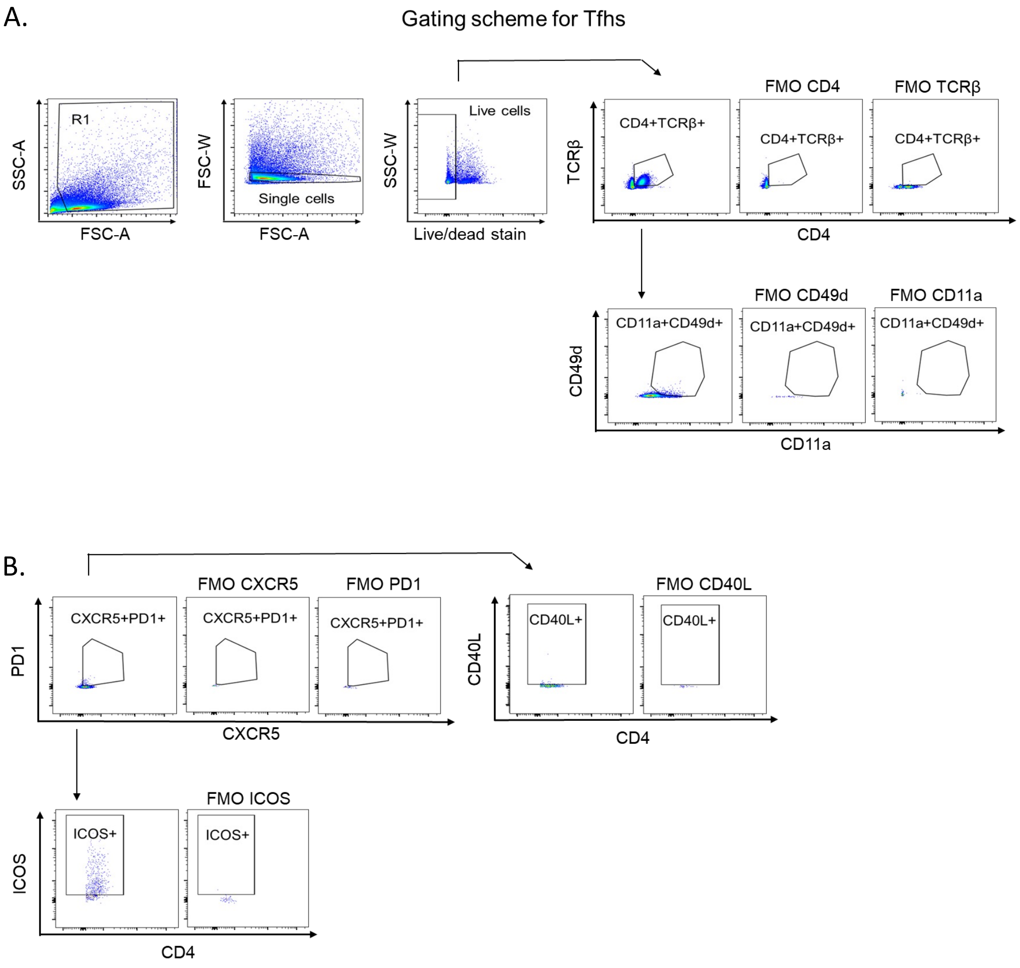


**Figure S1. Gating strategy for T follicular helper cells.** Adult C57BL/6 male and female mice were injected with PCV or PBS. Spleens and vaccine-draining lymph nodes were collected from all mice and assessed using flow cytometry. The gating strategy of the cells is shown. (A) Live single cells were gated on and the numbers of total CD4 T cells (CD4+TCRβ+) and antigen experienced (CD11a+CD49d+) CD4 T cells were determined. (B) Total and antigen experienced CD4 T cells were gated on and the number of Tfhs (CXCR5+PD1+) as well as expression of CD40L and ICOS on Tfhs were determined. Abbreviations: SSC-A = side scatter-peak area; FSC-A = forward scatter-peak area; FSC-W = forward scatter width; SSC-W = side scatter width; FMO = fluorescent minus one; TCR = T cell receptor.

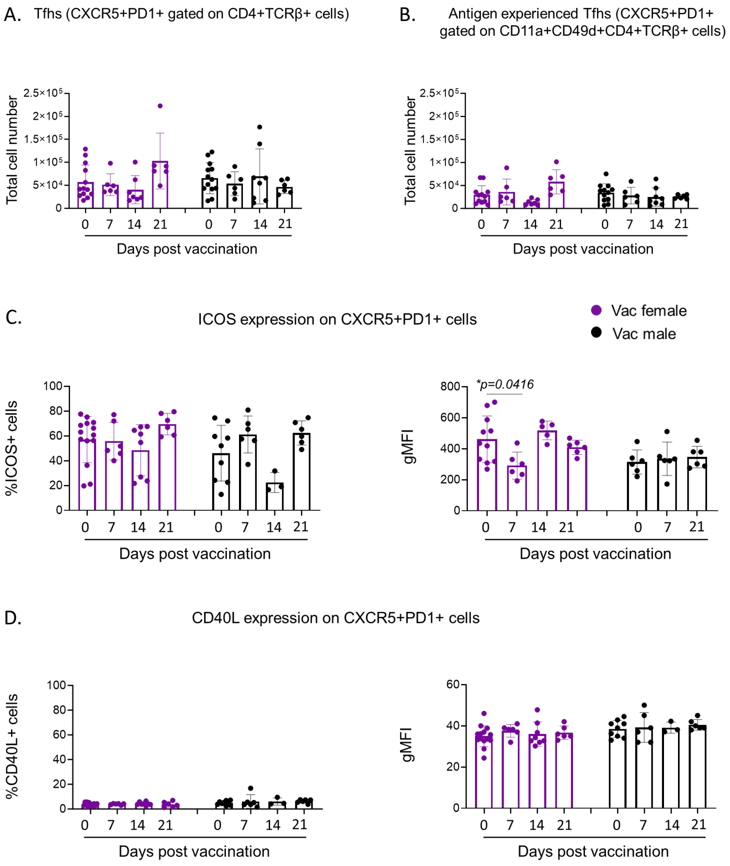


**Figure S2. No sex-based differences in T follicular helper cell responses in the spleen following PCV vaccination.** Adult C57BL/6 male and female mice were injected with PCV or PBS. Spleens were collected from all mice and assessed using flow cytometry. (A) Total and (B) antigen experienced Tfh cells were assessed at days 7, 14 and 21 post vaccination. Total Tfhs were assessed for their expression of (C) ICOS and (D) CD40L. (A) Total and (B) antigen experienced (CD11a+CD49d+) T cells (CD4+TCRβ+) cells were gated on and total numbers of CXCR5+PD1+ cells per mouse were determined. Total CXCR5+PD1+ cells were gated on and the percentage and geometric mean florescence intensity (gMFI) of (C) ICOS and (D) CD40L were determined. Data are pooled from 6 separate experiments and each dot represents an individual mouse. * denotes significant differences between the indicated groups as determined One-Way ANOVA followed by Šídák’s multiple comparisons test. Bar graphs represent the mean +/- SD.

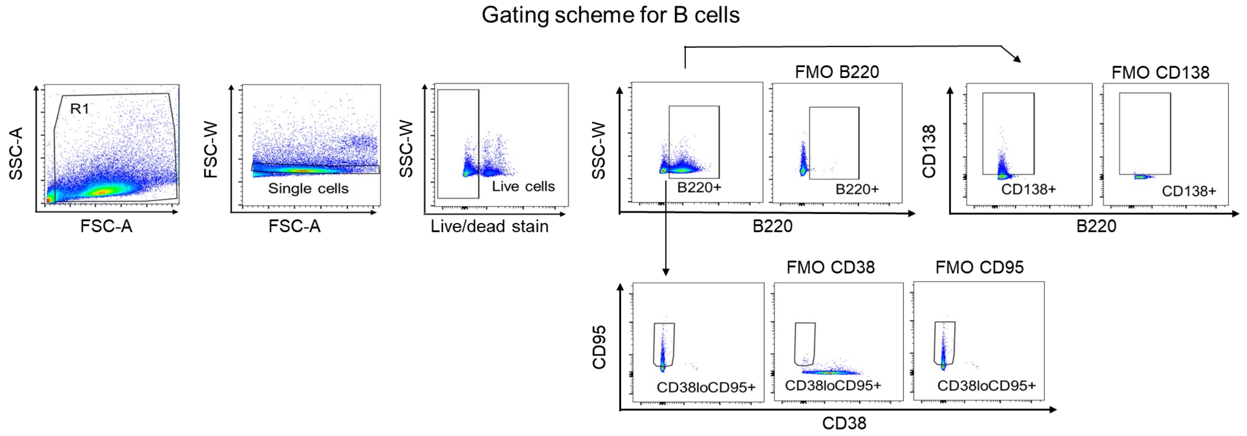


**Figure S3. Gating strategy for B cells.** Adult C57BL/6 male and female mice were injected with PCV or PBS. Spleens and LNs were collected from all mice and assessed using flow cytometry. The gating strategy of the cells is shown. Live single cells were gated on and the numbers of B cells (B220+) were determined. B220+ cells were gated on and the number of plasmablasts (CD138+) and GC B cells (CD38loCD95+) were determined. Abbreviations: SSC-A = side scatter-peak area; FSC-A = forward scatter-peak area; FSC-W = forward scatter width; SSC-W = side scatter width; FMO = fluorescent minus one.

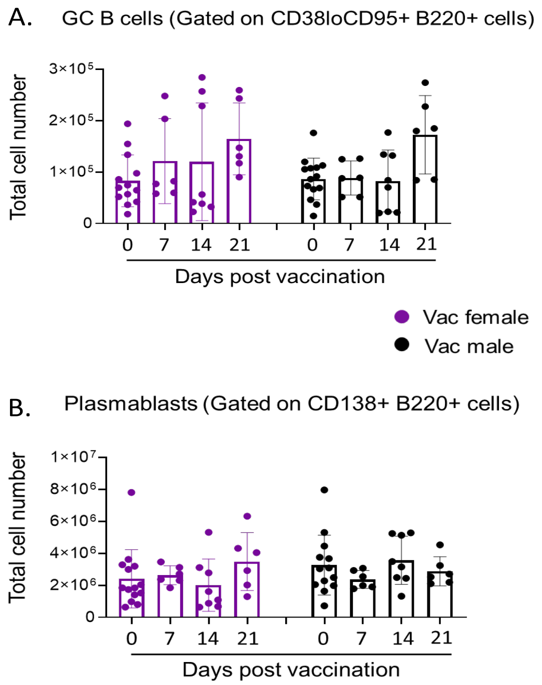


**Figure S4. No sex-based differences in B cell responses in the spleen following PCV vaccination.** Adult C57BL/6 male and female mice were injected with PCV or PBS. Spleens were collected from all mice and assessed using flow cytometry. (A) GC B cells and (B) plasmablasts were assessed at days 7, 14 and 21 post vaccination. Total B220+ B cells were gated on and total numbers of (A) GC B cells (CD38loCD95+) and (B) plasmablasts (CD138+) per mouse were determined. Data are pooled from 6 separate experiments and each dot represents an individual mouse. Bar graphs represent the mean +/- SD.

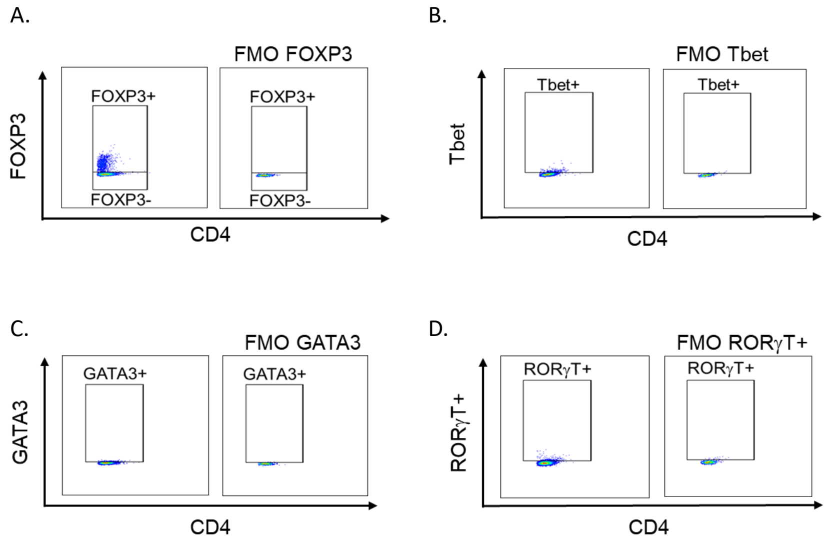


**Figure S5. Gating strategy for T cell subsets.** Adult C57BL/6 male and female mice were injected with PCV or PBS. Spleens and vaccine-draining lymph nodes were collected from all mice and assessed using flow cytometry. The gating strategy of the cells is shown. Live single antigen experienced (CD11a+CD49d+) CD4 T cells were gated on (as in Fig S1A) followed by gating on (A) Regulatory T cells ($FOXP3^{+}$), (B) Th1 ($Tbet^{+}FOXP3^{-}$), (C) Th2 ($GATA3^{+}FOXP3^{-}$) and (D) T helper 17 ($ROR\gamma T^{+}FOXP3^{-}$) subsets.

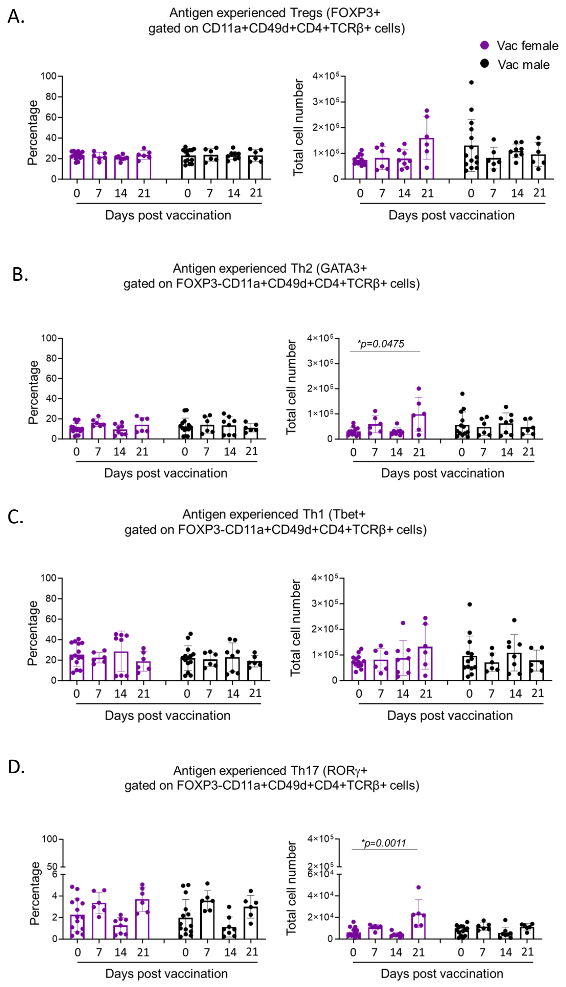


**Figure S6. No sex-based differences in T cell subsets in the spleen following PCV vaccination.** Adult C57BL/6 male and female mice were injected with PCV or PBS. Spleens were collected from all mice and assessed using flow cytometry. (A-D) Antigen experienced T cells ($CD11a^{+}CD49d^{+}CD4^{+}TCR\beta^{+}$) cells were gated on and percentages (left) and numbers (right) of T cell subsets per mouse were determined at days 7, 14 and 21 post vaccination. (A) Regulatory T cells ($FOXP3^{+}$), (B) Th2 ($GATA3^{+}FOXP3^{-}$), (C) Th1 ($Tbet^{+}FOXP3^{-}$) and (D) T helper 17 ($ROR\gamma T^{+}FOXP3^{-}$) were gated on. Data are pooled from 6 separate experiments and each dot represents an individual mouse. * denotes significant differences between the indicated groups as determined by (A&C) Kruskal-Wallis test followed by Dunn's multiple comparisons test or (B) One-Way ANOVA followed by Šídák's or Dunnett's multiple comparisons test. Bar graphs represent the mean +/- SD.

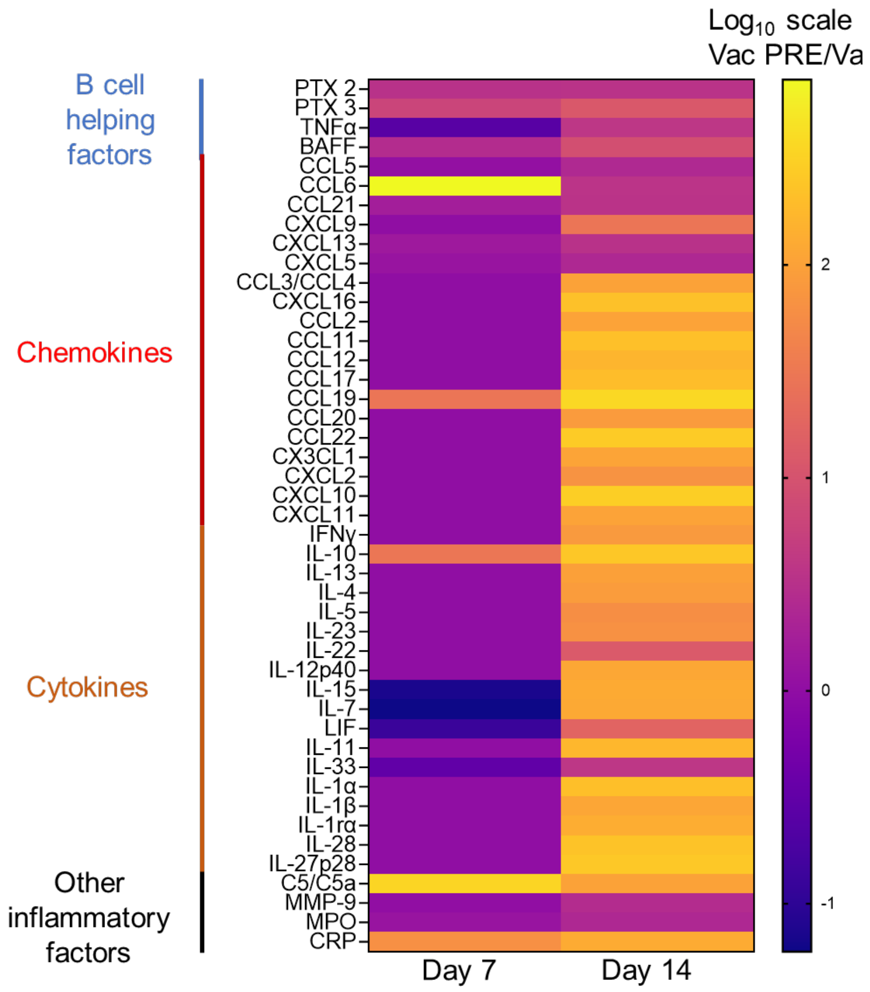


**Fig S7. Sex-based differences in the cytokine environment following PCV vaccination.** Adult C57BL/6 male and female mice were injected with PCV or PBS. Spleens were collected from all mice at days 7 and 14 post vaccination and assessed for cytokine/chemokines. Data are pooled from 3 separate experiments with 3 mice per group per timepoint and are presented in a heat map as $Log_{10}$ of the ratio of values from male mice divided by values from female mice.

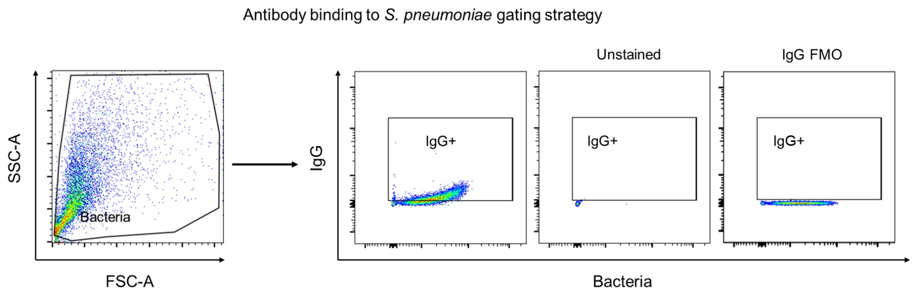


**Figure S8. Gating strategy for antibody binding to the surface of *S. pneumoniae*.** Adult C57BL/6 male and female mice were injected with PCV or PBS. Sera collected 4-weeks post vaccination were incubated with GFP-expressing *S. pneumoniae* to assess the binding efficacy of antibodies via flow cytometry. Following incubation with the sera, *S. pneumoniae* were washed and labeled with APC-tagged anti-mouse IgG. Bacteria (FITC+) were gated on and the percentage of IgG+ bacteria (FITC+APC+) were determined. Abbreviations: SSC-A = side scatter area; FSC-A = forward scatter area.

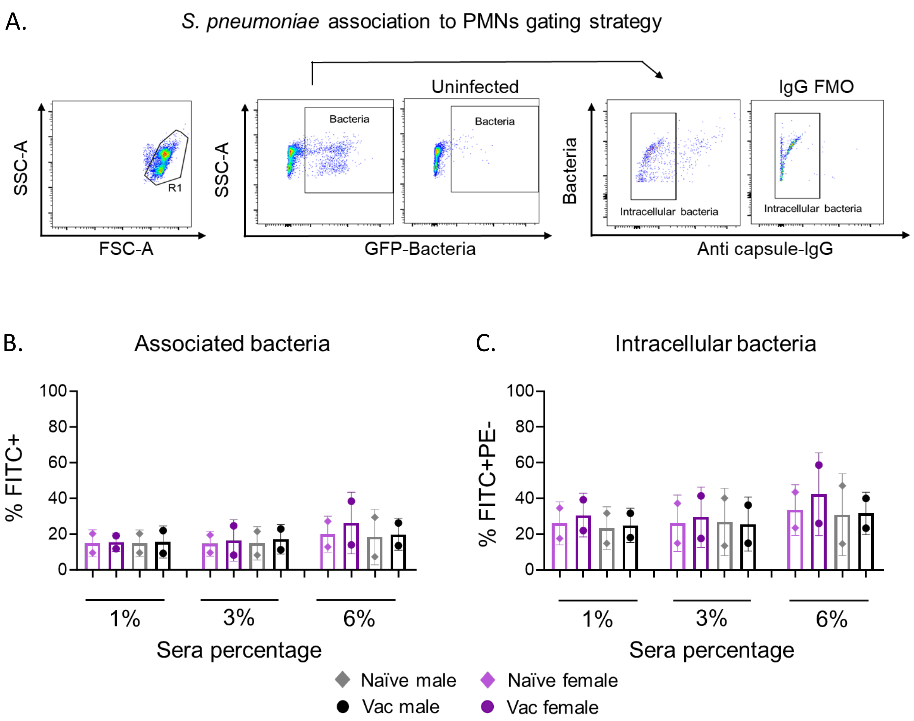


**Figure S9. No sex-based difference in the ability of sera from vaccinated mice to induce opsonophagocytic uptake of *S. pneumoniae* by PMNs.** Adult C57BL/6 male and female mice were injected with PCV or PBS. Sera collected 4-weeks post vaccination in all mice were used to opsonize GFP-expressing *S. pneumoniae* at 1-, 3- and 6% dilutions. Pre-opsonized GFP-expressing *S. pneumoniae* were incubated with PMNs and the reaction stained with PE-tagged anti-pneumococcal antibodies to measure intracellular vs extracellular bacteria via flow cytometry. (A) Gating strategy for bacterial uptake is shown. Bacteria (FITC+) were gated on and the percent of intracellular bacteria (FITC+PE-) was determined. (B) Percentages of associated (left) and intracellular (right) bacteria are shown. Data are pooled from 2 separate experiments with each dot representing the average of technical replicates per experiment. Bar graphs represent the mean +/- SD. Abbreviations: SSC-A = side scatter area; FSC-A = forward scatter area.

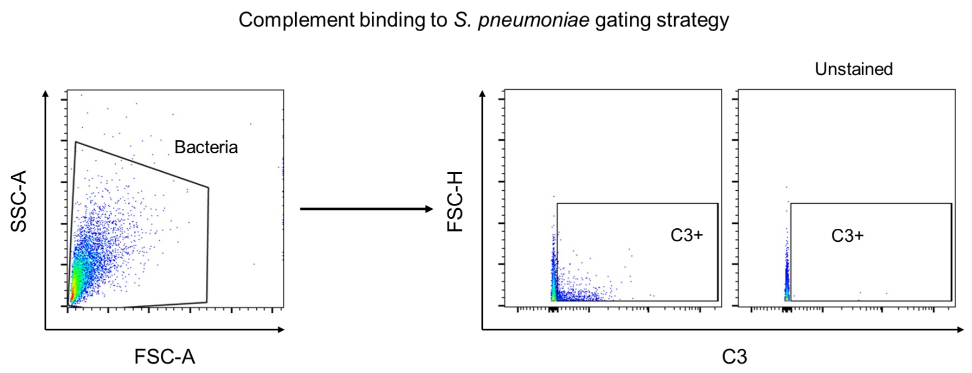


**Figure S10. Gating strategy for complement binding to the surface of *S. pneumoniae*.** Adult C57BL/6 male and female mice were injected with PCV or PBS. Sera collected 4-weeks post vaccination were incubated with *S. pneumoniae* to assess the binding efficacy of C3 via flow cytometry. Following incubation with the sera, *S. pneumoniae* were washed and labeled with FITC-tagged anti-C3 antibodies. Bacteria were gated on and the percentage of C3+ bacteria (FITC +) were determined. Abbreviations: SSC-A = side scatter area; FSC-A = forward scatter area.

| | | Average values | Standard deviation | P value (Significant?) |
|---|---|---|---|---|
| % antibody binding to *S. pneumoniae* | Vaccinated female 3% | 67.18 | 18.620 | 0.2482 (No) |
| | Vaccinated male 6% | 25.09 | 18.830 | |
| % opsonophagocytic killing with naïve PMNs | Vaccinated female 3% | 43.83 | 27.270 | > 0.9999 (No) |
| | Vaccinated male 6% | 26.34 | 23.180 | |
| % opsonophagocytic killing with matched vaccinated PMNs | Vaccinated female 3% | 37.32 | 28.570 | 0.3350 (No) |
| | Vaccinated male 6% | 67.06 | 5.924 | |
| % complement binding to *S. pneumoniae* | Vaccinated female 3% | 24.40 | 14.840 | 0.9944 (No) |
| | Vaccinated male 6% | 27.10 | 14.040 | |
| Fold change in binding to H292 cells | Vaccinated female 10% | 49.30 | 25.970 | > 0.9999 (No) |
| | Vaccinated male 20% | 39.44 | 18.0790 | |
| Fold change in bacterial growth | Vaccinated female 3% | 0.9911 | 0.4414 | 0.4382 (No) |
| | Vaccinated male 6% | 1.6310 | 0.2140 | |

**Table S1. Sex-based comparison in sera functions between matched antibody level conditions.** Adult C57BL/6 male and female mice were injected with PCV or PBS. Sera collected 4-weeks post vaccination in all mice was used to assess various antibody functions. This table summarizes the antibody functions assayed and compares conditions where the amount of antibody between vaccinated males and vaccinated females were equal.
